# Antagonism as a foraging strategy in microbial communities

**DOI:** 10.1101/2024.11.04.621785

**Authors:** Astrid K.M. Stubbusch, François J. Peaudecerf, Kang Soo Lee, Lucas Paoli, Julia Schwartzman, Roman Stocker, Marek Basler, Olga T. Schubert, Martin Ackermann, Cara Magnabosco, Glen G. D’Souza

## Abstract

In natural habitats, nutrient availability limits bacterial growth. We discovered that bacteria can overcome this limitation by acquiring nutrients through lysing neighboring cells via contact-dependent antagonism. Using single-cell live imaging and isotopic markers, we found that during starvation the type VI secretion system, which transfers toxins into neighboring cells, increases growth by enabling the uptake of nutrients from lysing cells. In a spatially structured environment, the gradual lysis of just a few target cells allows for the growth of a new antagonist cell. Genomic adaptations in antagonists, characterized by a reduced metabolic gene repertoire, and the prevalence of the type VI secretion system in natural environments, such as global oceans and soils, suggest that bacterial antagonism has the potential to substantially contribute to nutrient transfer within microbial communities across various ecosystems.

## Introduction

Bacteria require nutrients to synthesize cellular building blocks, such as nucleotides, amino acids and fatty acids, for growth. In nature, nutrients are often found only at low concentrations or in the form of complex polymers that require specific enzymatic degradation before they can be used as nutrients. Under such conditions, the biomass of other cells, rich in common metabolites and cellular building blocks, could serve as a valuable nutrient source. Killing neighboring cells can give access to these nutrients.

Bacteria have evolved various systems to lyse neighboring bacteria by transferring toxins, such as the type IV and type VI secretion systems (T4SS and T6SS) (*1*–*3*). These contact-dependent antagonistic systems are widespread in cultured bacteria (*2*). Their dominant ecological role is usually viewed as mediating competition for space and nutrients (*4*–*6*), leading to their characterization as ‘weapons’ of bacterial ‘warfare’ (*6, 2, 7*–*10*).

In this study, we ask whether contact-dependent antagonism enables bacteria to acquire nutrients from target cells, in addition to their role in removing competitors. This has been hypothesized previously (*11*), but, to the best of our knowledge, never been shown experimentally for non-predatory bacteria. Specifically, we focus on the widespread T6SS, a well-studied bacteriophage-like molecular machinery that enables transfer of diverse toxins into neighboring cells and that is present in approximately 25% of all sequenced Gram-negative bacteria (*12*).

## Results

### Contact-dependent antagonism enables growth during starvation

To determine whether the biomass of target cells provides a source of nutrients for antagonistic cells, we studied a simple community consisting of the marine isolates *Vibrio cyclitrophicus* ZF270 (target cells) and the T6SS-encoding *Vibrio ordalii* FS144 (T6SS cells) as an ecological model system. We investigated the growth of T6SS cells and the lysis of target cells at single-cell resolution in spatially structured communities growing in a microfluidic device (Fig. S1). When we provided the T6SS cells with a carbon source that they can metabolize (*N*-acetylglucosamine (GlcNAc), Fig. S2), the cells grew exponentially with a doubling rate of ∼0.2 h^-1^ (Fig. 1A, 1B, Fig. S3, Movie 1). In coculture with target cells, we observed a similar growth rate (Fig. S3, S5B). When we provided a carbon source that only target cells could metabolize (alginate, Fig. S2), T6SS cells failed to multiply in monoculture but were able to grow in coculture with target cells at a doubling rate of ∼0.07 h^-1^ (Fig. 1B, Fig. S3, Movie 2 and 3).

**Fig. 1:**
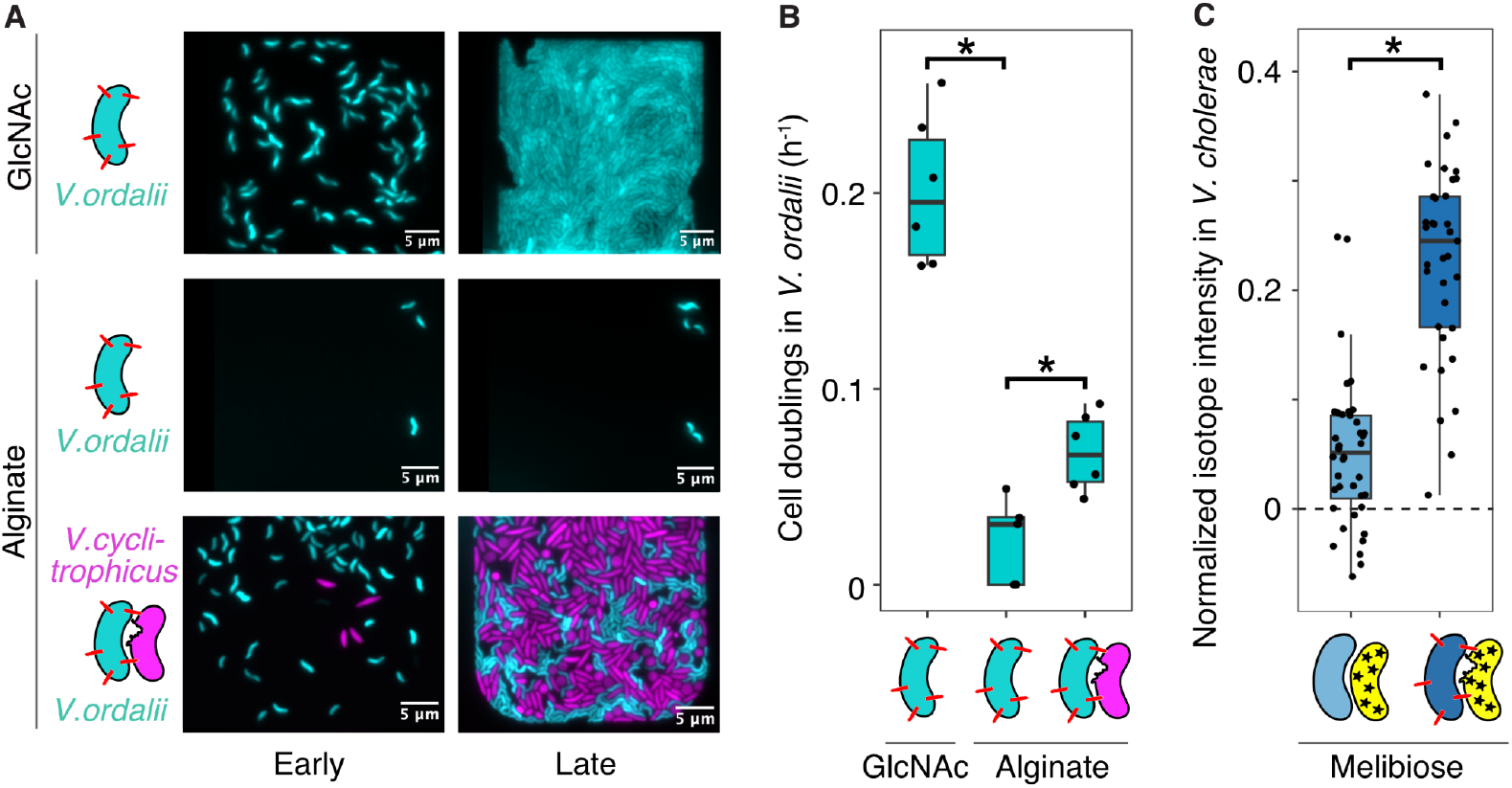
Growth benefit and nutrient acquisition through contact-dependent antagonism via the T6SS. (**A**) Representative images of T6SS cells (*V. ordalii* FS144, cyan) and target cells (*V. cyclitrophicus* ZF270, magenta) supplied with different carbon sources in microfluidic chambers at the start (0 h) and end time point (12 h, 16.7 h, and 24 h, top to bottom, respectively, due to different growth dynamics). (**B**) Growth rate of T6SS cells (*V. ordalii* FS144) in microfluidic chambers (n = 6, 5, 6; asymptotic Wilcoxon-Mann-Whitney test, *p =* 0.004 and 0.01, left to right). (**C**) Isotope incorporation by *V. cholerae* without T6SS (light blue) and with T6SS (dark blue) from deuterium-labeled *E. coli* (yellow). The deuterium signal in individual *V. cholerae* cells was quantified using Raman microspectroscopy. The coculture medium contained deuterium and melibiose as carbon source for continued growth of labeled *E. coli*. The values were normalized by subtracting the signal of the respective *V. cholerae* cells in this medium in monoculture (Fig. S5E). T6SS illustrated as red spikes, deuterium as star symbols (Wilcoxon rank-sum tests; T6SS-deletion vs. T6SS-encoding cells, *p* = 8 × 10^−13^).

To investigate whether the growth of T6SS cells without a metabolizable carbon source is facilitated by the T6SS-mediated killing of target cells, rather than the exchange of secreted nutrients (cross-feeding), we tested whether the growth is contingent on a functional T6SS. We used a genetic model system, namely *V. cholerae* 2740-80 and its T6SS-deletion mutant (*13, 14*), paired with *E. coli* as target cells. Again, we provided T6SS cells and target cells in microfluidic chambers with a carbon source that T6SS cells could not metabolize (melibiose, Fig. S1, Movie 4). We found that the growth rate of the T6SS-deletion mutant population was significantly lower than that of the T6SS wildtype population (Fig. S4 and S5A and C, Movie 5 and 6), suggesting that T6SS-mediated cell lysis increases the growth of T6SS cells under carbon starvation conditions.

To test whether the observed growth increase in T6SS cells is caused by the acquisition of nutrients from target cells, we used stable isotope probing (SIP)-Raman microspectroscopy (*15*). We first labeled *E. coli* target cells with deuterium and then cocultured them with unlabeled *V. cholerae* T6SS cells or the T6SS-deletion mutants. After 9 hours, T6SS cells showed significant levels of deuterium incorporation compared to the T6SS-deletion cells (Fig. 1C, Fig. S5D-E), further confirming that the T6SS is indeed mediating the uptake of nutrients.

### Slow cell lysis increases the nutrient uptake by surrounding cells

A more detailed examination of the T6SS-driven antagonism revealed that lysis is not instantaneous. We observed that the cell shape of attacked target cells changed from rod-shaped to round before bursting, likely due to T6SS effectors compromising cell wall integrity (*16*) (Table S2 and S3). These round *V. cyclitrophicus* target cells persisted for 1.43 ± 0.20 h in alginate medium and 0.33 ± 0.09 h in GlcNAc medium before bursting (mean ± 95% confidence interval) (Fig. 2A). Staining cells with propidium iodide (PI), which does not permeate intact cell membranes but can enter cells with leaky membranes (*17, 18*), we found that round target cells incorporated more PI than target cells in their usual rod shape (Fig. 2B). Together, these findings suggest that the action of carbon-starved T6SS cells causes slow lysis of target cells, likely leading to gradual rather than instantaneous nutrient release.

**Fig. 2:**
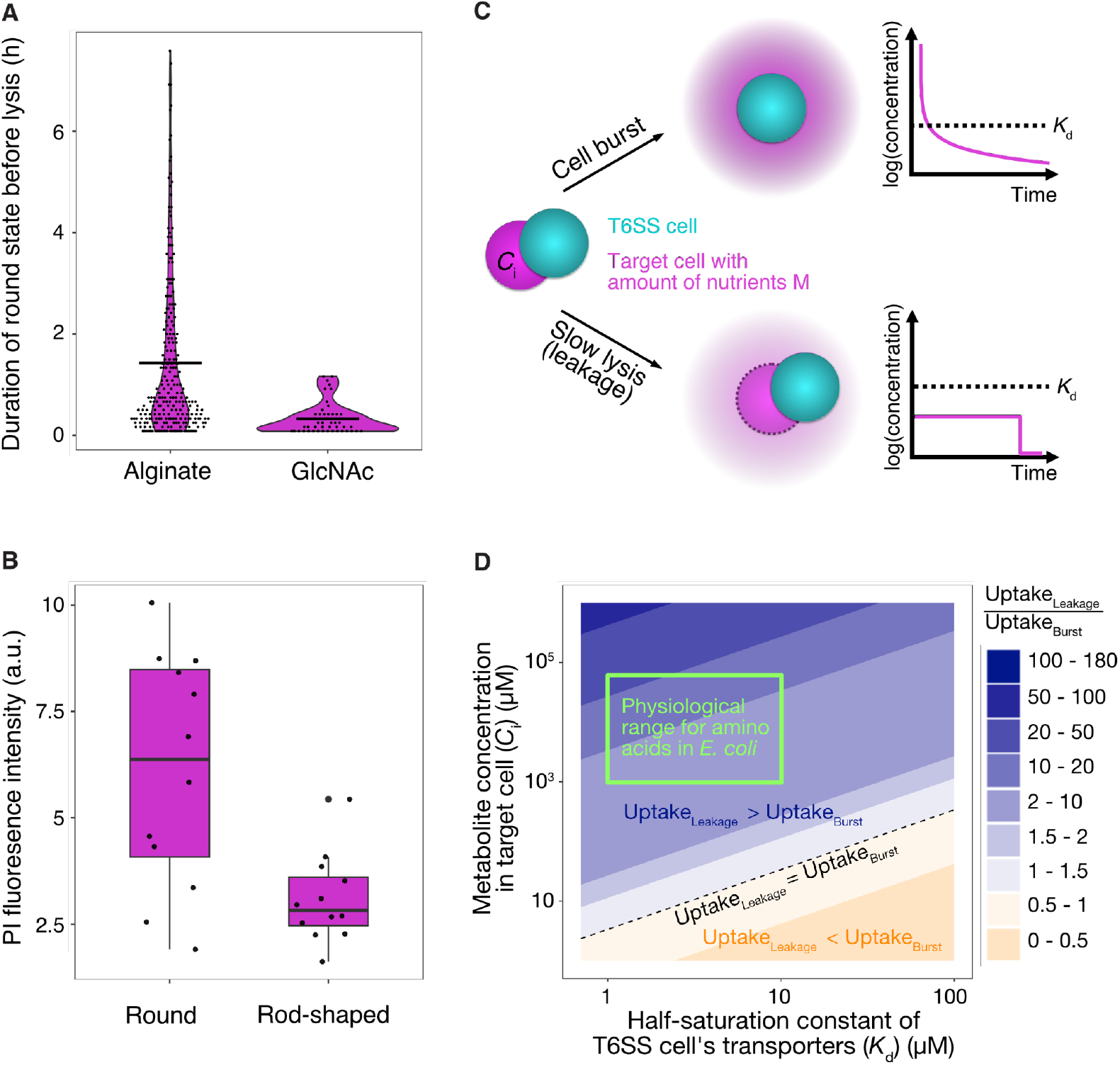
Slow cell lysis increases the nutrient gain for T6SS cells. (**A**) *V. ordalii* T6SS cells lyse the *V. cyclitrophicus* target cells slowly, especially under starvation conditions in alginate medium. (**B**) Induction of a round shape in target cells is associated with leakiness of the cell membrane, estimated by the propidium iodide (PI) staining intensity of round (length-to-width ratio of 1 to 1.8) and rod-shaped (length-to-width ratio of >1.8 to 6) target cells. Each dot represents the mean value from one microfluidic chamber (from 2 independent experiments with 6 chambers each). Jitter was applied in the horizontal direction. (**C**) Schematic diagram of the mathematical model that compares the nutrient concentration field available to T6SS cells through fast lysis (top) or slow lysis (bottom), alongside the nutrient release from the target cell over time (right). (**D**) Model predictions comparing the approximated total nutrient uptake by an antagonistic cell through slow and fast cell lysis, suggesting slow lysis increases nutrient uptake in the physiological parameter range. For slow lysis, we assume that the surface concentration of leaked metabolites is small compared to K_d_ of the T6SS cell. The green box depicts literature values for *C*_i_ and *K*_d_ in *E. coli* (Table S4).

To test whether slow cell lysis improves nutrient acquisition from target cells, we modeled the nutrient concentration field of a cell that bursts instantaneously in comparison with that of a cell that releases the same amount of nutrients slowly over time (Fig. 2C). We then modeled the nutrient uptake rate of a neighboring T6SS cell using Michaelis-Menten kinetics to take into account that membrane transport can reach saturation. Considering the typical intracellular concentration *C*_i_ of amino acids and the half-saturation constant *K*_d_ of amino acid transporters of *E. coli* (*19, 20*), we estimated that the approximate total nutrient gain of T6SS cells through slow lysis is 2-fold to 50-fold greater than that through fast lysis (Fig. 2C and D). Fast lysis led to larger initial concentrations of nutrients but exceeded the uptake capacity of the T6SS cell’s transporters, thus limiting its uptake. T6SS cells would acquire more nutrients from a bursting target cell than a slowly lysing target cell only if the target cell’s *C*_i_ was over 100-fold smaller or the *K*_d_ was over 10-fold higher than for amino acids in *E. coli*. These results suggest that T6SS cells gain more nutrients from slow cell lysis, which in our system was likely achieved by damaging cell membranes that then leak nutrients at concentrations that do not exceed the T6SS cell’s uptake capacity.

We found that the conversion of biomass from slowly lysing cells into new T6SS cells is very efficient. In the microfluidic chambers, on average one new T6SS cell formed for every 2 to 3 lysing target cells (Fig. S6B). This biomass conversion is substantially more efficient than previously reported for various taxa in well-mixed systems, which required 13 to 2,000 heat-killed cells for the growth of one new cell (*21*). This high efficiency may partly be explained by the about 2-fold larger volume of the target cells compared to the T6SS cells in our system (Fig. S6C to E). Nevertheless, this suggests that T6SS cells efficiently acquire nutrients from slowly lysing cells in spatially structured environments, potentially aided by sustained metabolite production of the lysing cells.

### Environmental significance of the employment of the T6SS for nutrient acquisition

If nutrient acquisition from cell lysate is a common use of the T6SS, one would expect genomic adaptations of T6SS-encoding bacteria toward a reduction of genes needed for the degradation of complex organic matter and synthesis of cellular building blocks, as observed in pathogenic and predatory bacteria (*22, 23*). To test this, we compared the gene repertoire of T6SS-encoding and T6SS-lacking genomes within the genus *Vibrio*. We analyzed ∼6,000 publicly available high-quality genomes. First, we screened the genomes for the presence of a T6SS (Fig. S7) and then de-replicated the genomes into 141 operational taxonomic units (OTUs) (Fig. S8). We found that 82 OTUs included only genomes with T6SS, 39 OTUs included only genomes without T6SS, and 20 OTUs included both T6SS-containing and T6SS-lacking genomes (Fig. S9). A pan-genome-wide association study (*24*) on the representative genomes of the exclusively T6SS-containing and T6SS-lacking OTUs was used to identify genes that were associated with the T6SS. As many bacterial traits are conserved among related taxa, we removed associations that arose from the underlying phylogenetic structure of the OTUs using a post hoc label-switching permutation test (*24*) (Fig. S10). We found 20 genes that had significantly positive and 157 genes that had significantly negative associations with the T6SS. The genes positively associated with the T6SS primarily encode the T6SS itself (Fig. 3A). The genes negatively associated with the T6SS are mainly involved in metabolic processes, such as carbohydrate metabolism, metabolite transport, amino acid metabolism, and biosynthesis of secondary metabolites (Fig. 3A). This suggests that these metabolic functions are reduced in species harboring a T6SS. As an example for carbohydrate metabolism, we tested whether T6SS-encoding *Vibrios* are commonly unable to degrade alginate, like in our experimental system. We found that indeed T6SS-encoding genomes contained significantly fewer enzymes to degrade alginate (1.54 ± 0.75 alginate lyases compared to 6.03 ± 2.24, disregarding phylogenetic structure) (Fig. 3B, Fig. S11E). The genome size and genome completeness were, however, not significantly different compared to T6SS-lacking OTUs (Fig. S11A-C). Overall, this reduced repertoire of metabolic genes, particularly for carbohydrate metabolism, in T6SS-encoding genomes is consistent with the expectation of genomic adaptation toward the acquisition of cellular building blocks from external sources, rather than their biosynthesis from simple metabolites obtained from the decomposition of complex organic matter.

**Fig. 3:**
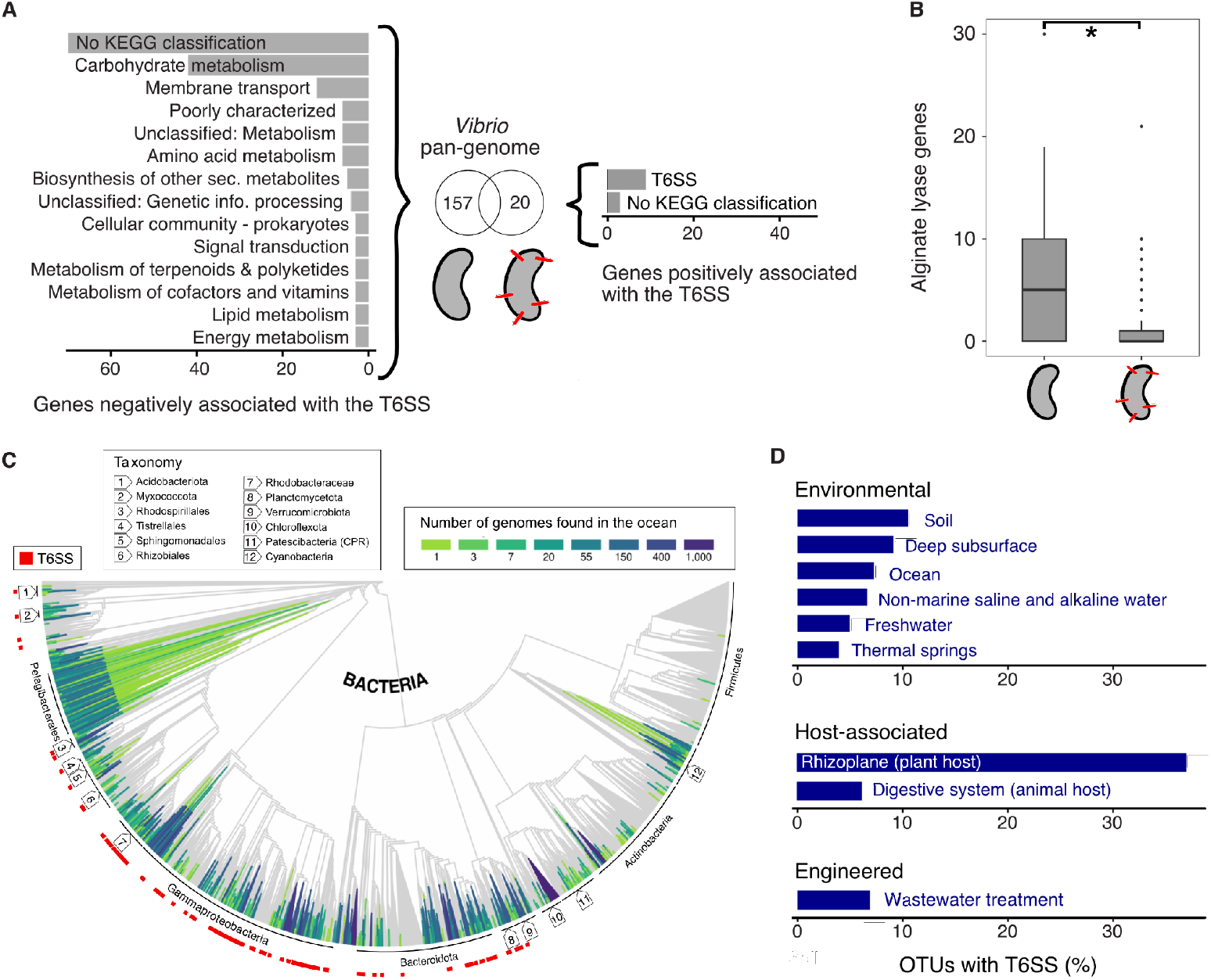
*Vibrio* with T6SS have a reduced metabolic gene repertoire and bacteria with T6SS are prevalent in natural environments. (**A**) Counts of genes that are positively or negatively associated with the presence of the T6SS in 141 *Vibrio* species-level operational taxonomic units (OTUs), grouped into functional categories according to KEGG classification. A gene can be part of several KEGG classes. Functional categories with less than 3 significant genes are not shown. (**B**) Number of alginate lyase genes across *Vibrio* OTUs with and without the T6SS. Wilcoxon rank sum test, *p* = 1×10^−5^. (**C**) Genomes of the Ocean Microbiomics Database (OMD) (green–blue) placed onto the bacterial Genome Taxonomy Database (GTDB) backbone tree (gray) (modified from Fig. 2C in Paoli et al. (*26*)) with red squares indicating T6SS-encoding genomes. For visualization, the last 15% of the nodes were collapsed (color codes indicate the number of genomes collapsed). (**D**) Prevalence of T6SS-encoding OTUs in different ecosystems based on data from the genomic catalog of Earth’s microbiomes (*28*).

To better understand the prevalence of T6SS across bacteria, we first focused on marine bacteria. In the ocean, cell lysis induced by viruses was identified as an important contributor to carbon and nutrient cycling, a phenomenon known as the ‘viral shunt’ (*25*). Likewise, cell lysis through bacterial antagonism may enhance this flux of carbon from living bacterial cells to lysis products. We screened the Ocean Microbiomics Database (OMD) (*26*), a consolidated collection of metagenome-assembled genomes from four major global ocean expeditions and two marine time series, as well as single-cell amplified genomes and marine microbial isolate genomes, for the presence of T6SS genes (Fig. S12). In the combined 7,610 bacterial species-level OTUs, we found the T6SS in 281 OTUs, i.e. ∼3.7%. In 229 of these OTUs more than half of the OTU members possessed a T6SS, i.e. ∼3.0% (Fig. 3C). Apart from clades that are known to be rich in T6SS, like *Gammaproteobacteria* and *Bacteroidota*, we also found that *Planctomycetota* and *Rhodobacteraceae*, which are known to be abundant in the coastal ocean (*27*), frequently encode the T6SS. Overall, the prevalence of the T6SS in marine taxa from the global oceans suggests that T6SS-mediated nutrient acquisition is common and may thus significantly influence nutrient cycling in the ocean.

A wider screen of natural environments showed that the T6SS is also common in diverse habitats. We screened for the presence of the T6SS in a genomic catalog of Earth’s microbiomes (*28*), containing 45,599 species-level OTUs that include metagenome-assembled bacterial genomes from over 10,000 samples from diverse habitats. We found the T6SS in 4–7% of OTUs in aquatic systems, 9–11% of OTUs in terrestrial systems, and 37% of OTUs in the rhizoplane (Fig. 3D). This is consistent with previous findings that suggest that T6SS-encoding bacteria are more common in structured than free-living habitats (*11*).

Also, soils have the largest average number of cells per volume of any known ecological environment (*29, 30*), and the rhizosphere is considered to be the richest niche within this habitat (*4, 31*), likely enhancing contact-dependent interactions. In the rhizoplane, most bacteria are thought to be commensals, but plant pathogens were found to use the T6SS to dominate the root microbiota and increase their virulence against the host plant (*4*). In environments that are comparatively rich in organic biomass, like the digestive systems of animals or wastewater treatment plants, 6% and 7% of OTUs were T6SS-encoding, respectively. These findings indicate that the T6SS is widespread across both nutrient-rich and nutrient-poor natural environments.

## Conclusion

In this work, we found that contact-dependent antagonism can serve to acquire nutrients. It allows bacteria to access the cell lysate, i.e., metabolites and cellular components, of neighboring cells as a nutrient source, which can be rapidly assimilated at low metabolic cost. In contrast to hard-to-digest complex organic compounds, such as polysaccharides, this source contains many common readily available cellular building blocks such as nucleotides from RNA and DNA, and amino acids from proteins (25% and 57% of cell content, respectively) (*32*). Polymerizing these building blocks into functional cell components represents an energetic saving of more than 90% in comparison with first having to synthesize them, and also brings a reduced proteomic cost (*32*). This way of nutrient acquisition is likely effective in microbial communities with ample cell–cell contact and limited diffusional loss of the freed nutrients, such as those in biofilms, soil aggregates, marine snow, the rhizosphere, and the intestinal mucus layer.

Nutrient acquisition is an alternative function to the mediation of bacterial competition and other functions which have been ascribed to the T6SS, such as acquisition of genetic material, kin discrimination, defense, and metal scavenging. These functions are not mutually exclusive, but the genomic signatures of T6SS-encoding species in *Vibrio* match the expectations that arise from a role in nutrient acquisition. The reduced repertoire of enzymes for carbohydrate degradation renders cells unable to grow on certain carbon sources, e.g. alginate in the case of *V. ordalii*. In these conditions, they are not in competition with other cells for the carbon source, but the employment of the T6SS enables them to acquire carbon from nearby cells. Under a broad definition of predation, i.e. killing cells for nutrient uptake, this interaction may be classified as facultative predation; however, there is no evidence that T6SS cells actively hunt target cells, and thus it does not meet stricter definitions of predation (*33*). In contrast to predation through ixotrophy, a recently described hunting strategy found in some filamentous T6SS-encoding bacterial species of the *Saprospiria* and *Cytophagia* classes with ‘grappling hooks’ to attach to prey cells (*34*), the T6SS cells studied here did not hold on to target cells.

As the T6SS is found in diverse taxa and diverse environments, we expect inter-bacterial cell lysis to also affect large-scale biogeochemical processes. Cell lysis releases both labile dissolved organic matter and carbon-rich particulate organic matter (*25*). The pool of labile substances of the lysate can be readily reused by bacteria (*35*) while the more recalcitrant particulate organic matter may preferentially sink into deeper ocean layers and thus contribute towards long-term carbon storage in the ocean, as proposed for the ‘viral shunt’ (*25*), caused by viral lysis of microorganisms. Our analyses suggest that there may be a parallel ‘bacterial shunt’ that participates in marine biogeochemical cycling.

## Acknowledgments

We thank current and previous members of the Schubert & Ackermann and Magnabosco labs and the PriME collaboration for helpful discussions, as well as Marco Gabrielli and Piersilvio De Bartolomeis. We thank Hans Wildschutte for providing the antibiotic deletion of *Vibrio ordalii* and Franziska Oschmann and Janis Fluri for the development of the segmentation and tracking tool *MIDAP*. We thank Knut Drescher and Niklas Netter for their advice on single cell segmentation of the T6SS cells. We thank Russell Naisbit for editorial comments and declare the use of ChatGPT-4 to improve the phrasing of individual sentences and to inspire specific coding solutions.

## Author contributions

Conceptualization: AS, MA, CM, GD

Methodology: AS, GD, FJP, KSL, JS, MB, LP

Investigation: AS, GD, FJP, KSL

Visualization: AS

Funding acquisition: MA, OS, GD, RS, JS

Project administration: AS

Supervision: MA, OS, CM, GD

Writing – original draft: AS

Writing – review & editing: AS, OS, MA, GD, CM, and all other co-authors

## Data and materials availability

All data and code will be available upon publication on ERIC Open (opendata.eawag.ch).

## List of Supplementary Materials

Materials and Methods

Figures S1 to S12

Tables S1 to S6

References

Movies S1 to S6

