## Supplemental material for "Antagonism as a foraging strategy in microbial communities"

**The PDF file includes**

Materials and Methods
Figures S1 to S12
Tables S1 to S6
References

**Further supplementary files**

Movies S1 to S6

### Materials and Methods

Growth media

For maintenance and pre-cultures, *Vibrio cyclitrophicus a*nd *Vibrio ordalii* were grown in Marine Broth (MB2216, DIFCO) whereas *Escherichia coli* and *Vibrio cholerae* strains were grown in Lysogeny Broth (LB, Merck) with the appropriate antibiotics (Table S1).

Experiments involving *V. cyclitrophicus* and *V. ordalii* were performed in Tibbles Rawling (TR) medium containing either 0.1% alginate or 0.1% N-acetyl-glucosamine (GlcNAc, weight/volume, Sigma-Aldrich) as described previously (*1*).

Experiments involving *E. coli* and *V. cholerae* were performed in M9 medium containing 0.1% melibiose or 0.1% glucose (weight/volume, Sigma-Aldrich) as described previously (*2*).

Bacterial strains

*V. cyclitrophicus* ZF270 and *V. ordalii* FS144 were obtained from an isolate collection derived from coastal waters, where isolates were obtained either from size-fractionated water samples, handpicked algal detritus particles and zooplankton, or different body parts of marine invertebrates (*1*). We used a mutant of *V. ordalii* that lacks an antibiotic production gene cluster (*3*) to rule out the possibility that antibiotic production was the antagonism mechanism involved in nutrient acquisition.

For all experiments described in this study, we used fluorescently tagged strains. Fluorescent *E. coli* (TB204 and TB205) was generated previously by genomically integrating a GFP or RFP gene, respectively (*4*). *V. cyclitrophicus* ZF270 had been tagged previously with a plasmid-based GFP marker (*5*). For this study, we tagged *V. ordalii* with the plasmid pVSV208, which carries the fluorescent dsRed marker gene. For this, we conjugated *V. ordalii* with *E. coli* DH5-alpha lambda pir carrying pVSV208 and *E. coli* DH5-alpha lambda pir carrying pEVS104. Briefly, 1 ml each of donor, recipient, and helper strain cultures grown for 16h in MB with the appropriate antibiotics were centrifuged at 12,000 rpm for 2 min and resuspended in 100 µl of antibiotic-free MB medium. Following this step, 100 µl of *E. coli* carrying pVSV208, 100 µl of *E. coli* carrying pEVS104, and 100 µl of recipient strains of interest were mixed. The mix of strains was centrifuged at 6000 g and the supernatant discarded. The cell pellet was then resuspended in 50 µl MB to make a thick slurry and spotted onto an MB agar plate. The plates were incubated overnight at room temperature (25 °C). The colonies on MB agar were resuspended in 50 µl MB and plated onto MB with 2.5 µg/mL chloramphenicol. These plates were incubated at room temperature for 24 h. Transconjugants were picked and screened for fluorescence and verified by Sanger sequencing.

*V. cholerae* 2740-80 T6SS-encoding strains (JS93 and B625) and the T6SS-deletion mutants *V. cholerae* 2740-80-Δ*hcp*1 Δ*hcp*2 (JS107 and AV009) were described previously (*6*, *7*). Descriptions and references of all genetic modifications are listed in Table S1.

Microfluidic experiments

The microfluidics experiments and microscopy were performed as described previously (*2*, *5*, *8*). Briefly, microfluidics experiments were performed in devices made of an inert polymer – polydimethylsiloxane (PDMS) – bound to a glass coverslip (Fig. S1). The PDMS layer had numerous growth chambers with dimensions of 30 x 30 x 0.88 µm (*l x b x h*). The medium flowed constantly through the main channel and diffused into the lateral growth chambers. Growth medium was supplied using 8-channel syringe pumps (New Era) at a flow rate of 0.1 ml $h^{-1}$. The sub-micron height of the growth chambers means that cells grow as a monolayer.

Automated microscopy imaging was performed using an inverted microscope (IX83, Olympus) driven by an automated stage controller (Marzhauser Wetzlar), shutter, and laser-based autofocus system (ZDC 2, Olympus). Many chambers of the same PDMS chip were imaged in parallel , and phase-contrast and fluorescent (GFP and/or TxRed) images of each position were taken every 5 min. The microscopy unit and PDMS chip was maintained at 25 °C or 37 °C (see below) using a cellVivo microscope incubation system (Pecon GmbH).

The *V. cyclitrophicus* and T6SS-encoding *V. ordalii* strains were pre-cultured in MB with 5 µg $ml^{-1}$ chloramphenicol for 16 h at 25 °C and 220 rpm, following which, 1 ml culture was centrifuged at 13,323 rpm for 2 min and the supernatant discarded. Cells were then resuspended in 1 ml TR medium with 0.1% alginate and incubated for 2 h at 25 °C for residual growth of the T6SS cells to occur (starvation phase). Cells were centrifuged at 12,000 rpm for 2 min and 950 µl of the supernatant discarded, then the cell pellet was resuspended in the remaining 50 µl medium. Target and T6SS-encoding or T6SS-deletion cells were mixed in a 10:2 ratio and 2 µl of this mix was pipetted into the microfluidic growth device. For *nutrient acquisition experiments*, devices were fed with TR medium containing 0.1% alginate, whereas for *competition experiments*, devices were fed with TR medium containing 0.1% *N*-acetyl-glucosamine (GlcNAc) at 25 °C. *E. coli* and *V. cholerae* strains were processed in a similar way but using LB medium for overnight cultures at 37 °C and 220 rpm. For *nutrient acquisition experiments*, devices were fed with M9 medium containing 0.1% melibiose, whereas for *competition experiments*, devices were fed with M9 medium containing 0.1% glucose at 37 °C.

For *nutrient acquisition experiments* (using alginate or melibiose, respectively, as carbon source), chambers containing mono- or cocultures were imaged for approximately 16 h (6 frames $h^{-1}$) followed by imaging at a higher frame rate (12 frames $h^{-1}$). In the case of *competition experiment*s (using GlcNAc or glucose, respectively, as carbon source), cells were processed similarly but without the 2 h starvation phase for the T6SS or T6SS-deletion mutant cells and imaging was conducted immediately with a high frame rate (12 frames $h^{-1}$). A table describing all imaged chambers with strains, conditions, and imaging times is available as Data 1.

Image visualization

Fluorescent channel images of representative chambers were displayed at initiation of growth, 0 h, and at a late timepoint, chosen based on the growth dynamics in the chamber and specified in the figure legends. The fluorescent signal from *V. cyclitrophicus* ZF270 cells was false-colored in magenta, *V. ordalii* FS144 cells in cyan, *E. coli* in yellow, and *V. cholerae* in blue using Fiji/ImageJ v2.9.0 (*9*).

Image analysis – segmentation and tracking of cells

Cell segmentation and tracking was performed using a custom image analysis pipeline MIDAP (v0.3 and higher, github.com/Microbial-Systems-Ecology/midap_public). Briefly, cell segmentation was performed using Omnipose (*10*) with the pre-trained model bact-fluor-omni-midap. Cell tracking was performed using STrack (*11*).

Image analysis – cell counts and duplication rate

To estimate the growth rates of *V. ordalii* and *V. cholerae* T6SS-deletion mutant cells, the following analysis was performed.

The number of cells within one chamber was computed for each image and for each strain, based on MIDAP's segmentation and tracking results of the respective fluorescent channel images. A generalized linear model was fitted to the number of cells over time for each chamber, assuming a logarithmic relationship, using the R function glm(cell number ~ time, family = poisson(link = 'log')). The raw data and the model fits are displayed in Fig. S3. The estimated slope of each fitted model was used to compute the doublings per hour. To this end, the doubling time in minutes was computed as $log_2$ / slope estimate. We then converted this doubling time in minutes to doublings per hour (i.e., the duplication rate).

The computed growth rate of chamber 3 of *V. ordalii* monoculture in alginate medium was removed from Fig. 1B. It was identified as an outlier by the boxplot analysis (ggplot::geom_boxplot function in R). We therefore investigated the raw imaging data and found that the cell count was increased through a cell entering the chamber from the main channel (after 1.5 h, frame 18) and its daughter cells. As this chamber initially contained only 1 resident cell, this led to a strong apparent increase in cell number.

The analysis was conducted using R software v4.1.2 in RStudio v2021.09.2+382.

Image analysis – area quantification in pixels

This analysis was performed to estimate the growth rate of the *V. cholerae* with intact T6SS. Since the T6SS cells have their T6SS sheath fused to GFP or mCherry (translational fusion reporter), the fluorescence of these cells manifests as foci and thus single-cell segmentation was not possible. We therefore used the pixel area of the fluorescent signal to estimate growth rates.

The total area of cells within one chamber was estimated for each image and for each strain based on the pixel count with fluorescent signal over a threshold of 50. A generalized linear model was fitted to the area of cells over time for each chamber, assuming a logarithmic relationship, using the R function glm(area in µm2 ~ time, family = poisson(link = 'log')). The raw data and the model fits are displayed in Fig. S4. The estimated slope of each fitted model was used to estimate the doublings of area in μ$m^2$ per hour. To this end, the doubling time was computed as $\log_2$ / slope estimate. We then converted this area doubling time to area doublings per hour by computing 1/(area doubling time). The analysis was conducted using R software v4.1.2 in RStudio v2021.09.2+382.

Persistence time of antagonized cells

We identified target cells that were already attacked by the T6SS by their round cell shape. Target cells with a length-to-width ratio between 1 and 1.8 were considered to have a round cell shape. Target cells with a length-to-width ratio of 1.8 or more were considered to be rod-shaped. The persistence time of the round cells was computed as the time from which a target cell turned from rod-shaped to round until the cell disappeared from the chamber. Only cells that stayed round until they disappeared were considered. The analysis was conducted using R software v4.1.2 in RStudio v2021.09.2+382.

Stable isotope probing–Raman microspectroscopy

To label target *E. coli* cells (TB205) with deuterium ($D_2O$ from Silantes), *E. coli* target cells tagged with RFP were grown for 24 h in 100% $D_2O$-containing M9 minimal medium supplemented with 0.1% melibiose. Corresponding unlabeled *E. coli* target cells were grown for 24 h in 100% $H_2O$-containing M9 minimal medium supplemented with 0.1% melibiose. *V. cholerae* B625 and *V. cholerae* T6SS-deletion mutant (AV009) cells were grown for 16 h in LB medium. These were centrifuged at 13,323 rpm for 2 min, supernatants discarded and cell pellets washed twice with M9 salts minimal medium without a carbon source. The cell pellet was then incubated for 9 h in M9 salts minimal medium without a carbon source to terminate any residual metabolic activity. Deuterated and non-deuterated *E. coli* cultures (2 ml) were also centrifuged at 12,000 rpm for 2 min but cell pellets were resuspended in 100% $D_2O$-containing M9 minimal medium supplemented with 0.1% melibiose. 200 μl labeled or 200 μl unlabeled target cells were incubated with either T6SS-encoding or T6SS-deletion cells in 100% $D_2O$-containing M9 medium supplemented with 0.1% melibiose. As controls, T6SS-encoding or T6SS-deletion strains were incubated alone in either deuterated or non-deuterated M9 minimal medium. These cultures were incubated for 9 h at 37 °C without shaking.

To measure individual cells (T6SS, T6SS-deletion, and *E. coli*) using Raman microspectroscopy, 1.5 μl droplets of each sample were placed on an aluminum-coated slide (EMF Corp., USA). The samples on the slide were dried in a 30 °C incubator for 15 min, rinsed with Milli-Q water to remove traces of the medium, and air-blown to dry. This process enabled the individual cells to be distributed on the aluminum-coated slide for single-cell Raman measurement.

A commercial confocal Raman microspectroscope (LabRAM HR Evolution, Horiba Scientific, France) was used for the measurements. This system is based on an upright microscope (BxFM, Olympus) integrated with optical components for Raman measurement, including a Raman laser (continuous wave neodymium-doped yttrium aluminum garnet – CW Nd:YAG; 532 nm with 10 mW), an objective (MPlan N 100×, 0.90 NA, Olympus), a grating (300 grooves/mm; blazed at

600 nm), and a detector (back-illuminated deep depleted CCD). Individual cells were measured within the spectral window of 300–3,400 $cm^{-1}$, which covers most cellular signals, with a 10-sec exposure time of the Raman laser and a 100-µm confocal pinhole.

To differentiate between the T6SS and target cells, the cytochrome *c* peak at 750 $cm^{-1}$ was used: *V. cholerae* exhibited this peak, whereas *E. coli* did not (Fig. S5D). The raw Raman data were processed using a code built in-house (*12*). Specifically, for the spectral window of 1,800–3,200 $cm^{-1}$, the data underwent smoothing (de-noising) and baseline subtraction using the Savitzky–Golay filter (with polynomial order 3 and frame length 7) and a polynomial-based algorithm (2nd order polynomial), respectively. To evaluate the deuterium labeling status of the cells, the ratio CD / (CD + CH) = $I_{2,040–2,300}$ / ($I_{2,040–2,300}$ + $I_{2,800–3,100}$) was calculated, where *I* represented integrated intensity.

Propidium iodide staining

Cocultures of *V. cyclitrophicus* and *V. ordalii* cells in microfluidic growth devices were propagated as described above. Following 30 h of propagation, cells were stained with propidium iodide (PI, Thermo Fisher Scientific) by injecting 2 µl dye into microfluidic channels from the inlet ports. Cells were then incubated for 1 h and then imaged using GFP and TxRed filters. To determine the staining intensities, target cells were segmented using the GFP channel and cellular properties such as length-to-width ratio and the intensity normalized to area in the TxRed channel (indicating PI intensity) were computed for each cell. The analysis was conducted using R software v4.1.2 in RStudio v2021.09.2+382.

Mathematical modeling of nutrient uptake from cell bursts or slow cell lysis

*Nutrient release by cell burst*

The release of nutrients by a single pulse from a point source modeling the burst of an individual target cell is described in 3D by

$$C(r,t) = \frac{M}{\sqrt{4\pi D t}^3}\, exp\left(-\frac{r^2}{4\, D\, t}\right) \qquad (1)$$

with $C(r, t)$ being the concentration of nutrients at a distance *r* from the point source at a given time *t* after initial release. Here, *M* is the total amount of nutrients released by the pulse, and *D* is the diffusion coefficient of the nutrient released.

Note that this expression is an approximation of the lysis patch released by a cell containing a mass *M* of nutrient, due to the consideration of a point source instead of a spatially extended initial source. A simple way to recover some spatial extension of the source at $t = 0$ is to approximate the initial patch by a Gaussian patch of typical radius *a,* with *a* the known radius of the cell before lysis (a typical approach, see for example Clerc et al. (*13*) and references therein).

To do so, we determine the time $t_s$ at which the Gaussian patch given by equation (1) has a standard deviation in space equal to *a*, that is

$$t_s = \frac{a^2}{2D}. \qquad (2)$$

One can then use the new pulse expression

$$C(r,t) = \frac{M}{\sqrt{4\pi D(t_s + t)}^{\,3}} \, exp\,(-\frac{r^2}{4D(t_s + t)}) \qquad (3)$$

for $t \geq 0$, with now non-infinite concentration and non-zero spatial extent at $t = 0$. Note that for a typical diffusivity $D \approx 1 \times 10^{-9}$ m$^2$ s$^{-1}$ and typical cell radius $a \approx 1$ µm, we have $t_s \approx 5 \times 10^{-4}$ s, which is extremely small compared to the typical minute timescale considered for uptake. This regularization of the initial patch thus has negligible impact on the results.

*Nutrient release by leakage*

For the case of leakage from a slowly lysing cell, we consider the release of nutrients from a spherical source of radius *a* with a release rate per unit exposed surface area φ (in moles per area per time). Assuming steady-state of the concentration profile around the source in 3D, we have

$$C(r,t) = \frac{a}{r}\frac{\phi a}{D}$$

that is, $C(r,t) = \frac{\phi a^2}{Dr}$ (4)

where as before, $C(r, t)$ is the concentration of nutrients at a distance $r$ from the center of the source at time $t$. As before, $a$ is the radius of the leaking target cell, and $D$ is the diffusion coefficient of the leaked nutrients.

The concentration of nutrients in the immediate neighborhood of the leaking cell (i.e., $r = a$) is thus given by

$$C(r = a) = \frac{\phi a}{D}. \qquad (5)$$

Considering a total amount of nutrients $M$ that can be released from the leaking cell (assuming no replenishment of the resource, and steady release rate $\phi$ as a first approximation), the duration of nutrient leakage until the nutrient source is exhausted is given by

$$t_{max} = \frac{M}{4\pi a^2 \phi}. \qquad (6)$$

*Nutrient uptake*

The nutrient uptake by a neighboring T6SS cell is assumed to follow Michaelis–Menten kinetics, which captures both the linear increase of uptake rate with surrounding nutrient concentration increase for low concentrations, and the saturation of uptake rate at large concentration due to the finite speed and number of a cell's transporters. We thus consider an uptake rate per cell given by

$$u = V_{max}\frac{C}{C + K_d} \qquad (7)$$

with $V_{max}$ the maximal uptake rate (at saturation of the uptake capability of the cell), $C$ the local nutrient concentration, and $K_d$ the half-saturation constant (i.e., the nutrient concentration corresponding to half-maximal uptake rate).

Note that we make an approximation by not modeling how the cell uptake modifies the surrounding nutrient patch.

*Nutrient uptake through cell burst*

The total nutrient uptake of a cell exposed to a nutrient pulse (in total moles) can be obtained by direct integration in time of the instantaneous uptake rate in the burst conditions, that is

$$U_{tot}^{pulse} = \int_0^\infty u(C^{burst}(t))dt \qquad (8)$$

$$= \int_0^\infty V_{max} \frac{C^{burst}(t)}{C^{burst}(t) + K_d} dt \qquad (9)$$

$$= V_{max} \int_0^\infty \frac{1}{1 + \frac{K_d}{M} \sqrt{4\pi D t}^{\,3}} dt \qquad (10)$$

where we used equation (1) taken at $r = 0$ to obtain the concentration profile in the burst case $C^{burst}(t)$. We thus assume that a T6SS cell is exposed to the concentration at the center of the lysis patch.

By defining the new integration variable $\tau = (\frac{K_d}{M})^{\frac{1}{3}} \sqrt{4\pi D t}$, we obtain

$$U_{tot}^{pulse} = \frac{V_{max}}{2\pi D} (\frac{M}{K_d})^{\frac{2}{3}} \int_0^\infty \frac{\tau}{1 + \tau^3} d\tau \quad . \qquad (11)$$

Using the fact that

$$\int_0^\infty \frac{\tau}{1 + \tau^3} d\tau = \frac{2\pi}{3\sqrt{3}}, \qquad (12)$$

we obtain the total uptake

$$U_{tot}^{pulse} = \frac{V_{max}}{3\sqrt{3}D} (\frac{M}{K_d})^{\frac{2}{3}} \quad . \qquad (13)$$

*Nutrient uptake through nutrient leakage*

Combining the known concentration in the immediate neighborhood of the leaking cell given by equation (5) and the Michaelis–Menten expression of uptake rate from equation (7), the nutrient uptake rate $u$ of a cell in that scenario is

$$u = V_{max} \frac{\frac{\phi a}{D}}{\frac{\phi a}{D} + K_d} .$$

The total nutrient uptake $U_{tot}$ by a cell in the immediate neighborhood of the leaking cell is thus obtained by integrating this uptake rate over the total leakage duration $t_{max}$ (equation (6)), that is

$$U_{tot}^{leak} = \frac{V_{max}}{\phi a + K_d D} \times \frac{M}{4\pi a} . \qquad (14)$$

*Nutrient uptake through cell leakage vs. cell burst*

The ratio of the total nutrient uptake from a leaking cell $U_{tot}^{leak}$ (equation (14)) and a bursting cell $U_{tot}^{pulse}$ (equation (13)) is given by

$$\delta = \frac{U_{tot}^{leak}}{U_{tot}^{pulse}}$$

$$= \frac{3\sqrt{3}}{4\pi} \frac{1}{a} \left(\frac{M}{K_d}\right)^{\frac{1}{3}} \frac{1}{1 + \frac{\phi a}{DK_d}}$$

$$= \frac{3\sqrt{3}}{4\pi a} \left(\frac{M}{K_d}\right)^{\frac{1}{3}} \frac{1}{1 + \frac{C_s}{K_d}} \quad (15)$$

with $C_s$ being the concentration at the surface of the leaky cell, given by $C_s = \frac{\phi a}{D}$. We note $\eta = \frac{\phi a}{DK_d} = \frac{C_s}{K_d}$ the ratio of the concentration at the surface of a leaking cell to the half-saturation of uptake, and $\delta' = \frac{3\sqrt{3}}{4\pi a} \left(\frac{M}{K_d}\right)^{\frac{1}{3}}$. We thus have

$$\delta = \frac{\delta'}{(1+\eta)} \quad . \quad (16)$$

If $\eta \ll 1$, this expression can be further approximated linearly in $\eta$ by

$$\delta = \delta' (1 - \eta). \quad (17)$$

In that case, $\delta'$ captures most of the uptake advantage of leakage over burst. Considering an internal concentration $C_i$ of nutrient inside the source cell, where the total mass of nutrient $M$ inside the source cell can be described by $M = 4\pi a^3 \frac{C_i}{3}$ we observe that

$$\delta' = \frac{3\sqrt{3}}{4\pi a} \left(\frac{M}{K_d}\right)^{\frac{1}{3}}$$

$$= \frac{3\sqrt{3}}{4\pi} \left(\frac{4\pi}{3} \frac{C_i}{K_d}\right)^{\frac{1}{3}}$$

$$= \frac{3^{7/6}}{(4\pi)^{2/3}} \left(\frac{C_i}{K_d}\right)^{\frac{1}{3}} \quad (18)$$

Since for most nutrient compounds, the internal concentration $C_i$ is much larger than the half-saturation constant of uptake $K_d$, we will have $C_i \gg K_d$ and thus $\delta' > 1$, which means that the leakage strategy will be advantageous over the cell burst strategy.

This is visualized in the phase diagram in Fig. 2D, where the dashed line separates the parameter space where nutrient leakage results in more nutrient uptake (blue area) and the parameter space where a nutrient burst results in more nutrient uptake (orange area). The green square marks the literature values of the $K_d$ of amino acid transporters and $C_i$ for amino acids in *E. coli* at 37 °C (*14*, *15*). Note that Fig. 2D uses the expression of $\delta'$ given in equation (18), since we assume that generally $\eta << 1$, as we consider the surface concentration of leaked metabolites to be small compared to uptake constant $K_d$ in the case of slow leakage of a cell. Indeed, considering a large internal concentration $C_i = 60\,000$ $\mu$M and a small $K_d = 1$ $\mu$M together with a radius $a = 0.5$ $\mu$m and a diffusivity $D = 1000$ $\mu m^2/s$ (*16*), and estimating the typical leakage time $t_{max} \sim 1$ h (Fig. 2A), we can estimate that $\eta \sim 0.001$ which validates this approximation. Smaller values of $C_i$, larger values of $K_d$, or longer values of $t_{max}$ would result in even smaller estimates of $\eta$.

<u>Cell volume estimation</u>

To compare the volume of the target and T6SS cells, we computed the approximate cell volume based on segmented cell length and width measurements. These were obtained from cocultured target and T6SS cells in microfluidic chambers grown on alginate. Cell segmentation was performed using a custom image analysis pipeline MIDAP v3, as described above. For both target and T6SS cells, we computed the mean length and width of cells in the segmented images, averaging over the cells from four microfluidic chambers in the time between 16.6 and 24 hours (Fig. S6). We then used these average values to approximate cell volume. Given the approximately cylindrical shape of both cell types, we estimated cell volume $V$ using the formula for a cylinder $V = \pi \ x \ (\frac{width}{2})^2 \ x \ length$ . The estimated average cell volume of target cells was $V_{target} = \pi \ x \ (\frac{0.68 \ \mu m}{2})^2 \ x \ 1.90 \ \mu m = 0.69 \ \mu m^3$ and the estimated average cell volume of T6SS cells was $V_{T6SS} = \pi \ x \ (\frac{0.54 \ \mu m}{2})^2 \ x \ 1.61 \ \mu m = 0.37 \ \mu m^3$ .

The relative volume difference between target and T6SS cells was then calculated by comparing their mean volumes: $\frac{V_{target}}{V_{T6SS}} = \frac{0.69 \ \mu m^3}{0.37 \ \mu m^3} = 1.86$. Target cells were about 2-fold larger in size than T6SS cells. All analyses were conducted using R software v4.1.2 in RStudio v2021.09.2+382.

<u>Pangenomic association study</u>

For the pangenomic association study, we downloaded high-quality *Vibrio* genomes from RefSeq with the search term 'Vibrio.*'. We obtained 6478 genomes (downloaded on February 23, 2023, at 16:01). We removed all files containing 'ViralProj', as they result from sequencing of *Vibrio* phage. We also filtered the genomes for a genome completeness of ≥95% estimated by CheckM (*17*). The remaining 6062 genomes were used for the following analysis.

We screened all genomes for the genes of the T6SS subtype i (T6SSi), the subtype which is found in proteobacteria such as *Vibrio*, using the profile Hidden Markov Models (HMMs) published by Abby et al. (*18*) and the software HMMER v3.1b2 (hmmer.org) with an e-value threshold of 1e-5. The number of T6SS genes and the presence of each T6SS gene are shown in Fig. S3. We considered genomes to be T6SS-encoding when more than ⅔ of the 14 essential genes for the T6SSi were present in a genome, i.e., 10 or more, based on Abby et al. (*18*)).

We then compared the similarity of these *Vibrio* genomes by pairwise comparison, using the fastANI algorithm to compute their average nucleotide identity (ANI) and coverage (i.e., the part that can be aligned to each other) between all genomes using the software dRep v3.2.2 (*19*) (Fig. S4). We observed a gap in the ANI distribution of the *Vibrio* genomes at ca. 94% (Fig. S4), and therefore used a threshold of 94% ANI to dereplicate the genomes into clusters of OTUs. We obtained 141 OTU clusters. One genome from cluster 0_12 and 2 genomes from cluster 0_73 were removed, as they were part of very large and otherwise T6SS-encoding clusters and the T6SS was likely missing due to the incompleteness of these three genomes. Specifically, genome GCF_026613815.1_ASM2661381v1 is part of cluster 0_12 with 1676 T6SS-encoding genomes, but we found only 29% of the essential T6SS genes in this genome. The genomes GCF_024744975.1_ASM2474497v1 and GCF_026162605.1_ASM2616260v1are part of cluster 0_73 with 2040 T6SS-encoding genomes, but we found only 64% and 21% of essential T6SS genes, respectively. As a result, we continued our analysis with 82 clusters that contained only

T6SS-encoding genomes, 39 clusters that contained only T6SS-lacking genomes, and 20 clusters of genomes that contained both T6SS-encoding and T6SS-lacking genomes.

The best representative genome for each OTU was chosen by the software dRep. We screened the representative genomes of the 141 OTUs for the presence of carbohydrate-active enzymes, i.e., enzymes cataloged in the database of Carbohydrate-Active enZYmes (CAZy), using the HMMs published on dbCAN v2 (*20*).

We next used the software Roary v3.13.0 (*21*) to retrieve the pangenome from the 141 cluster representatives. Here, all protein sequences were clustered by 90% amino acid similarity, resulting in a pangenome of 273,727 protein-encoding genes. Of these, 162 were present in 99% to 100% of the genomes ('core genes'), 97 were present in 95% to 99% of the genomes ('soft core genes'), 2,214 were present in 15% to 95% of the genomes ('shell genes'), and the majority, 271,254 genes, were present in less than 15% of the genomes ('cloud genes').

We then estimated the phylogenetic structure of the 141 OTU representatives. We used the GTDBtk v2.3.2 to infer a phylogenetic tree (*22*). In brief, multiple sequence alignments were formed from the concatenation of 120 phylogenetically informative marker genes (bac120). The phylogenetic tree was then inferred with FastTree v2.1.10 (*23*) from the concatenated alignment of the 120 bacterial marker genes. We carried out this analysis with all three substitution models that are implemented in FastTree to describe amino acid evolution, namely the WAG (Whelan-Goldman), LG (Le-Gascuel), and JTT (Jones-Taylor-Thornton) substitution models. The three resulting phylogenetic trees were visualized using iTOL v6.9 (*24*). We added the bootstrap values as color, displayed the taxonomic annotation using GTDBtk v2.3.2, and denoted T6SS-encoding clusters, T6SS-lacking clusters, and mixed clusters with strain-specific presence of the T6SS.

To evaluate the association of a gene with either T6SS-encoding or T6SS-lacking genomes while accounting for phylogenetic relationships of the genomes, we used the software Scoary v1 (*25*). We used the software to score the presence of the T6SS in a genome for associations to a binary phenotype (i.e., the presence of another gene) using Fisher's test and accounted for population structure using a post hoc label-switching permutation test. This post hoc permutation test is based on the pairwise comparisons algorithm, alleviating the need to set mutation rate parameters (*26*). It computes the empirical $p$-value for each gene as $p = (r + 1)/(N + 1)$, with $r$ being the number of test statistics observed to be higher or equal to the unpermuted statistic and $N$ being the number of permutations. Genes with an empirical $p$-value of 0.05 or less (based on either of the three phylogenetic trees) were considered significantly associated with the T6SS, either positively or negatively.

We then grouped the genes that were positively or negatively associated with the T6SS into broad functional KEGG categories. For this, we used the software BlastKOALA v2.3 (*27*) to annotate all genes with KEGG's *K* numbers. Next, we added the higher-level classifications of KEGG to each annotated *K* number. The gene counts for each KEGG category were summed. KEGG categories with 3 or more significant genes are displayed in Fig. 4A. (We used the 'level B' of the KEGG categories to summarize the negatively T6SS-associated genes. For the positively T6SS-associated genes, the T6SS genes constituted KEGG categories with 3 or more significant genes and were therefore simply denoted as T6SS in Fig. 4A.)

Screen of the Ocean Microbiomics Database (OMD) for the T6SS

The Ocean Microbiomics Database (OMD) was used as a metagenomic dataset (*28*). This dataset included metagenomes from samples collected by Tara Oceans (*29*, *30*) (virus-enriched, $n$ = 190; and prokaryote-enriched, $n$ = 180), BioGEOTRACES expeditions ($n$ = 480), the Hawaiian Ocean Time-series (HOT, $n$ = 68), the Bermuda-Atlantic Time-series Study (BATS, n = 62) (*31*) and the Malaspina expedition ($n$ = 58) (*32*). The retrieved metagenome-assembled genomes (MAGs) were combined with 830 manually curated MAGs (*33*), 5,969 single-amplified genomes (SAGs) (*34*) and 1,707 REFs and 682 SAGs (*35*) of marine bacteria and archaea into a combined collection of 34,799 genomes.

All protein sequences that were comprised in the OMD (genes-redundant.faa) were screened with profile Hidden Markov Models (HMMs) for the T6SS subtype i (T6SSi), T6SS subtype ii (T6SSii), and T6SS subtype iii (T6SSiii) (*18*) using the software HMMER v3.1b2 with an e-value of $10^{-5}$. We further quality-filtered the HMMER hits based on the protein length, requiring the protein hit to be similar in length to the T6SS HMM (≥70% and ≤130% of the HMM).

GTDB backbone with OMD

All genomes in the OMD detected across 1,038 ocean metagenomes were placed onto the GTDB backbone tree (*36*) to reveal the extent of the phylogenomic coverage of the OMD (*28*). Clades without any genome in the OMD are colored gray. For visualization, the last 15% of the nodes were collapsed (modified from Paoli et al. (*28*)).

To compute the fraction of T6SS-encoding genomes, we first identified the T6SS genes in each of the 34,799 genomes, based on our screen of the redundant genes of the OMD. Then, genomes with more than ⅔ of the number of essential genes for the T6SS were considered T6SS-encoding (10 or more for the T6SSi, 12 or more for the T6SSii, and 8 or more for the T6SSiii, based on Abby et al. (*18*)). The 34,799 genomes were clustered using a 95% whole-genome average nucleotide identity cut-off into 8,304 prokaryotic species-level clusters or operational taxonomic units (OTUs) (*28*), with 7,610 bacterial clusters. We computed the fraction of T6SS-encoding genomes for each 'species cluster'. These fractions were displayed as barplots on the phylogenetic tree in Fig. 3C. The backbone was adopted from Paoli et al. (*28*), and shows the bacterial tree of life based on GTDB with the number of genomes included in the OMD.

Screening the Genomic Catalogue of Earth's Microbiomes (GEMs) collection for the T6SS

The genome collection from 'A Genomic Catalogue of Earth's Microbiomes' (GEMs) is available at https://portal.nersc.gov/GEM/ (*37*). This genome collection includes over 10,000 metagenomes collected from diverse habitats covering all of Earth's continents and oceans, human- and animal-host associated microbiomes, engineered environments, and natural and agricultural soils. It contains 52,515 metagenome-assembled genomes (MAGs) representing 12,556 novel candidate species-level operational taxonomic units (OTUs), spanning 135 phyla.

These 52,515 MAGs as well as 524,046 reference genomes had been clustered into 45,599 species-level OTUs. All genomes in this collection of OTUs had been evaluated using CheckM and were required to have a quality score >50 and were taxonomically annotated using GTDB-tk. Representative genomes had been selected with the highest CheckM quality scores with isolate genomes prioritized over MAGs and SAGs (*37*).

We screened the representative genomes of all OTUs for the presence of essential T6SS genes with the HHM for T6SSi, T6SSii, and T6SSiii (*18*) using the software HMMER v3.1b2 with an e-value threshold of $10^{-5}$. We considered genomes T6SS-encoding when more than ⅔ of the number of essential genes for the T6SS were present in a genome (10 or more for the T6SSi, 12 or more for the T6SSii, and 8 or more for the T6SSiii, respectively). We grouped the OTUs by their associated ecosystem type. In Figure 3B, we present the fraction of T6SS-encoding OTU representatives for each ecosystem, including only ecosystem types with 50 or more OTUs and meaningful ecosystem categories (excluding, e.g., ‘city’ or ‘defined media’) and merging the two wastewater plant categories ‘Nutrient removal’ and ‘Anaerobic digestor sludge’.

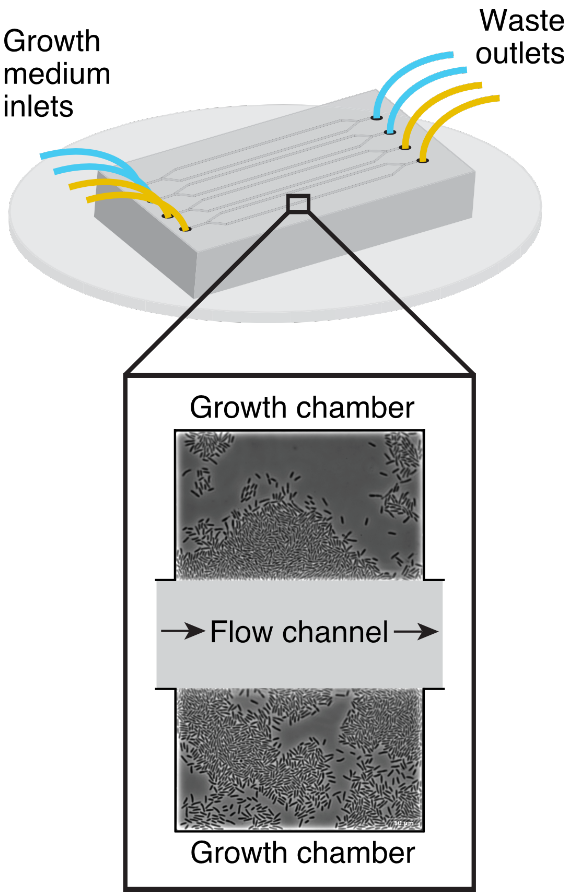


**Fig. S1. Schematic diagram of microfluidic system.**
Growth medium is provided through the inlet, flowing through the main flow channel and leaving through the outlet. Along the flow channel are multiple growth chambers, which allow cells to grow in monolayers. Different media (indicated by orange and blue) can be provided to growth chambers through different flow channels. Adapted from Stubbusch et al. (*38*).

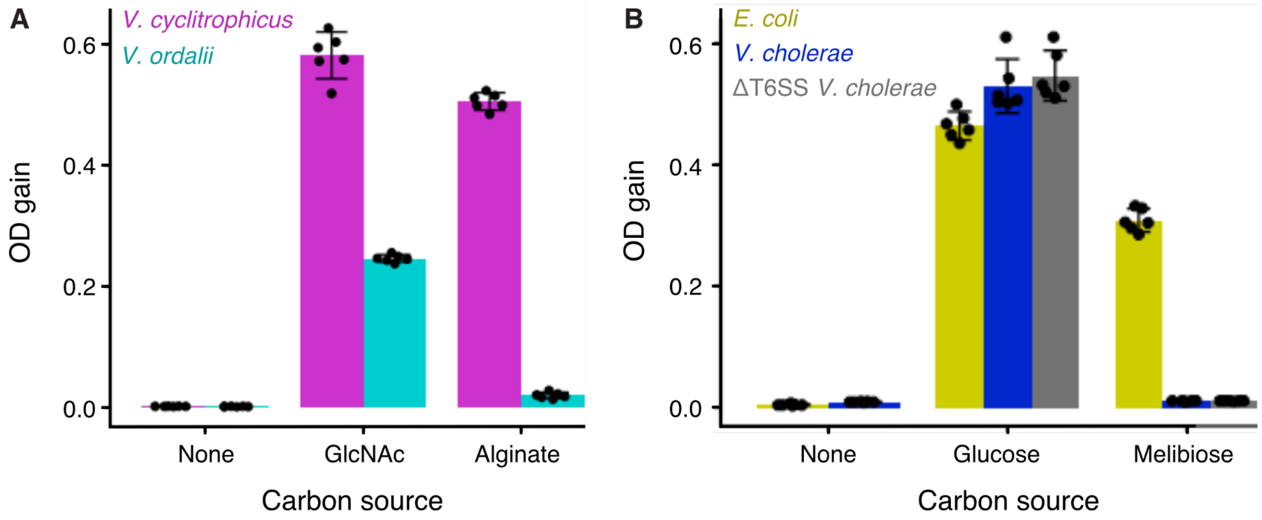


**Fig. S2. Growth yield of bacterial monocultures on different carbon sources quantified by optical density (OD).**
(A) OD gain of *V. cyclitrophicus* ZF270 (magenta) and *V. ordalii* FS144 (cyan) in monoculture in growth medium without carbon source (none), growth medium with *N*-acetylglucosamine (GlcNAc), a glucose derivative that constitutes bacterial cell walls, and growth medium with alginate, a polysaccharide from brown algae. (B) OD gain of *E. coli* MG1655 (yellow), *V. cholerae* 2740-80 (blue), and *V. cholerae* 2740-80 T6SS-deletion (gray) in monoculture in growth medium without carbon source (none), growth medium with glucose, and growth medium with melibiose, a disaccharide consisting of galactose and glucose. Error bars indicate 95% confidence intervals.

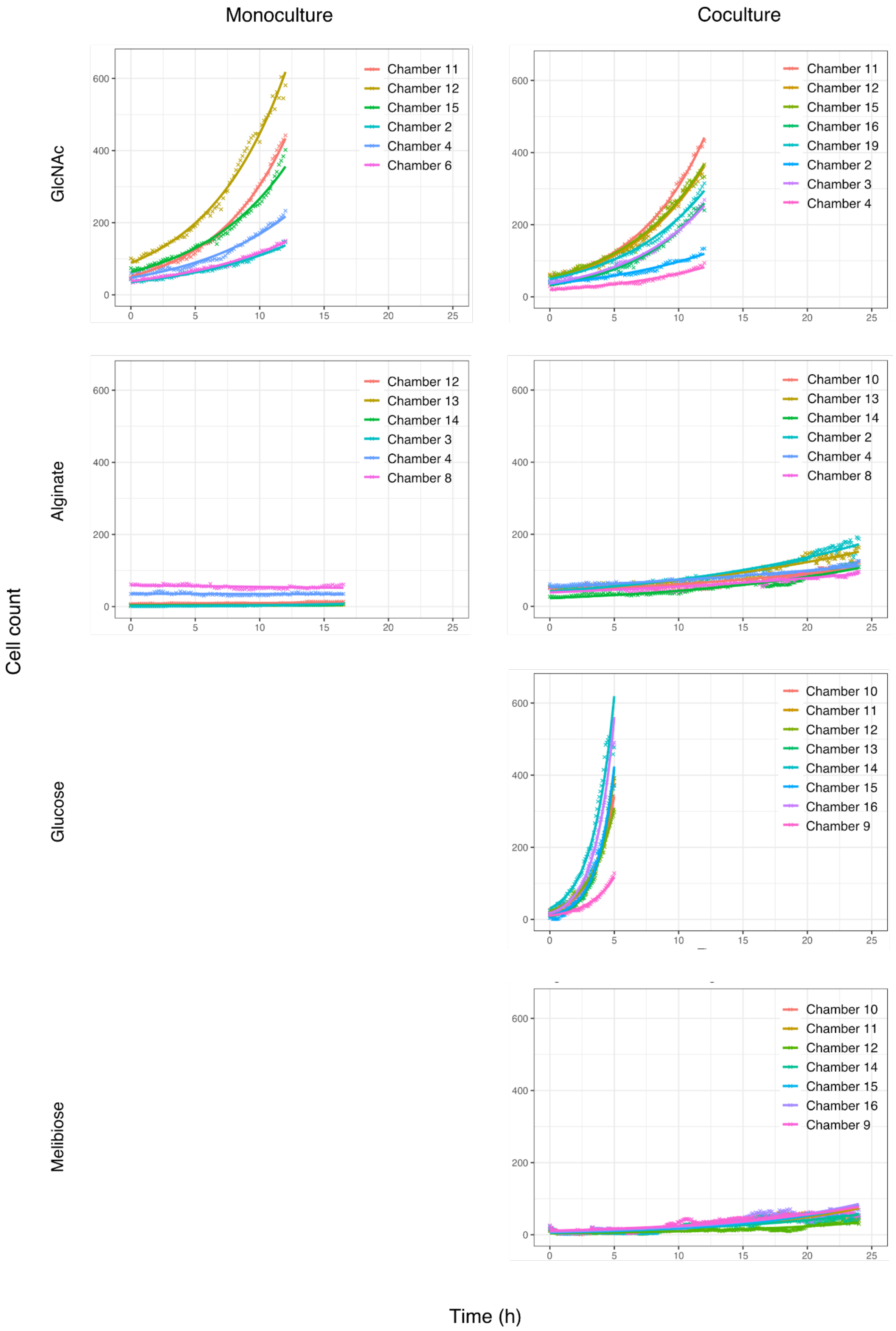
Monoculture
Coculture
GlcNAc
Alginate
Glucose
Melibiose
Cell count
Time (h)
Chamber 11
Chamber 12
Chamber 15
Chamber 2
Chamber 4
Chamber 6
Chamber 11
Chamber 12
Chamber 15
Chamber 16
Chamber 19
Chamber 2
Chamber 3
Chamber 4
Chamber 12
Chamber 13
Chamber 14
Chamber 3
Chamber 4
Chamber 8
Chamber 10
Chamber 13
Chamber 14
Chamber 2
Chamber 4
Chamber 8
Chamber 10
Chamber 11
Chamber 12
Chamber 13
Chamber 14
Chamber 15
Chamber 16
Chamber 9
Chamber 10
Chamber 11
Chamber 12
Chamber 14
Chamber 15
Chamber 16
Chamber 9

**Fig. S3. Cell numbers of *V. ordalii* in mono- and coculture subjected to different carbon sources.**

Cell count as a function of time in monocultures of *V. ordalii* and cocultures of *V. ordalii* and *V. cyclitrophicus* in microfluidic chambers provided with various carbon sources (see Materials and Methods). The number of *V. ordalii* cells reflects the number of segmented *V. ordalii* cells. Cell numbers are shown for each chamber individually (colors). Lines show fits of an exponential function, i.e., log(cell count) ~ intercept + slope x time. The estimated slope of these exponential fits was used to compute the doubling rate displayed in Fig. 1B.

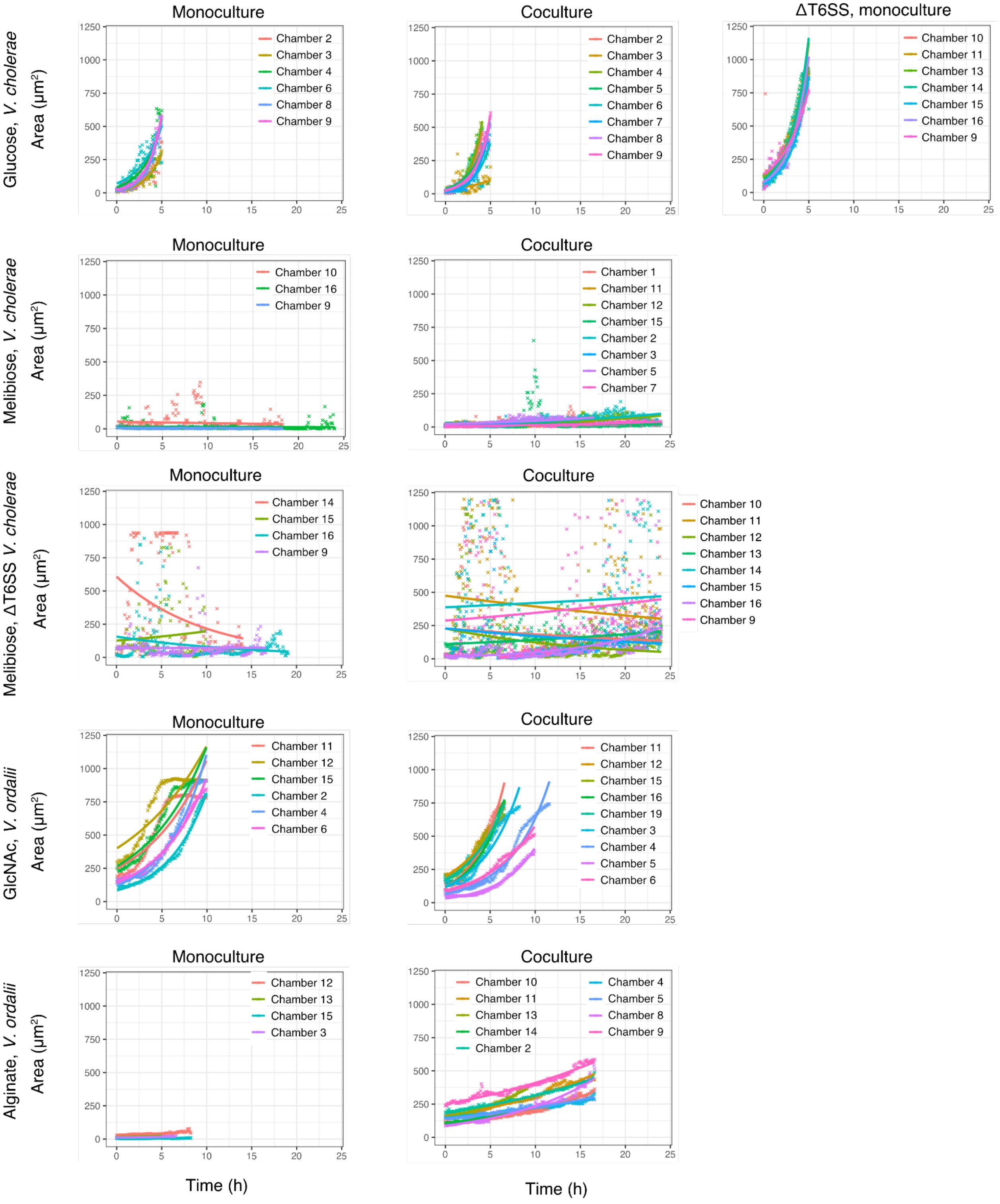
Monoculture
Coculture
ΔT6SS, monoculture
Glucose, V. cholerae
Area (µm²)
Chamber 2
Chamber 3
Chamber 4
Chamber 6
Chamber 8
Chamber 9
Chamber 2
Chamber 3
Chamber 4
Chamber 5
Chamber 6
Chamber 7
Chamber 8
Chamber 9
Chamber 10
Chamber 11
Chamber 13
Chamber 14
Chamber 15
Chamber 16
Chamber 9
Melibiose, V. cholerae
Area (µm²)
Chamber 10
Chamber 16
Chamber 9
Chamber 1
Chamber 11
Chamber 12
Chamber 15
Chamber 2
Chamber 3
Chamber 5
Chamber 7
Melibiose, ΔT6SS V. cholerae
Area (µm²)
Chamber 14
Chamber 15
Chamber 16
Chamber 9
Chamber 10
Chamber 11
Chamber 12
Chamber 13
Chamber 14
Chamber 15
Chamber 16
Chamber 9
GlcNAc, V. ordalii
Area (µm²)
Chamber 11
Chamber 12
Chamber 15
Chamber 2
Chamber 4
Chamber 6
Chamber 11
Chamber 12
Chamber 15
Chamber 16
Chamber 19
Chamber 3
Chamber 4
Chamber 5
Chamber 6
Alginate, V. ordalii
Area (µm²)
Chamber 12
Chamber 13
Chamber 15
Chamber 3
Chamber 10
Chamber 11
Chamber 13
Chamber 14
Chamber 2
Chamber 4
Chamber 5
Chamber 8
Chamber 9
0
250
500
750
1000
1250
0
5
10
15
20
25
Time (h)
Time (h)

**Fig. S4. Cell growth of *V. ordalii* and *V. cholerae* as population area over time.** Monocultures of *V. ordalii* and *V. cholerae* and cocultures of *V. ordalii* with *V. cyclitrophicus* and *V. cholerae* with *E. coli* were grown in microfluidic chambers provided with different carbon sources, namely glucose and melibiose for *V. cholerae* and GlcNAc and alginate for *V. ordalii* (see Materials and Methods). The area of the T6SS cells over time was computed, as it was not possible to obtain single-cell numbers through the segmentation of the cells in the case of *V. cholerae* with functional T6SS. Cell areas are shown for each chamber individually (colors). Lines show fits of an exponential function, i.e., log(cell area) ~ intercept + slope x time. The estimated slope of these exponential fits was used to compute the doubling rate of one $\mu m^2$ of cell area displayed in Fig. S5D.

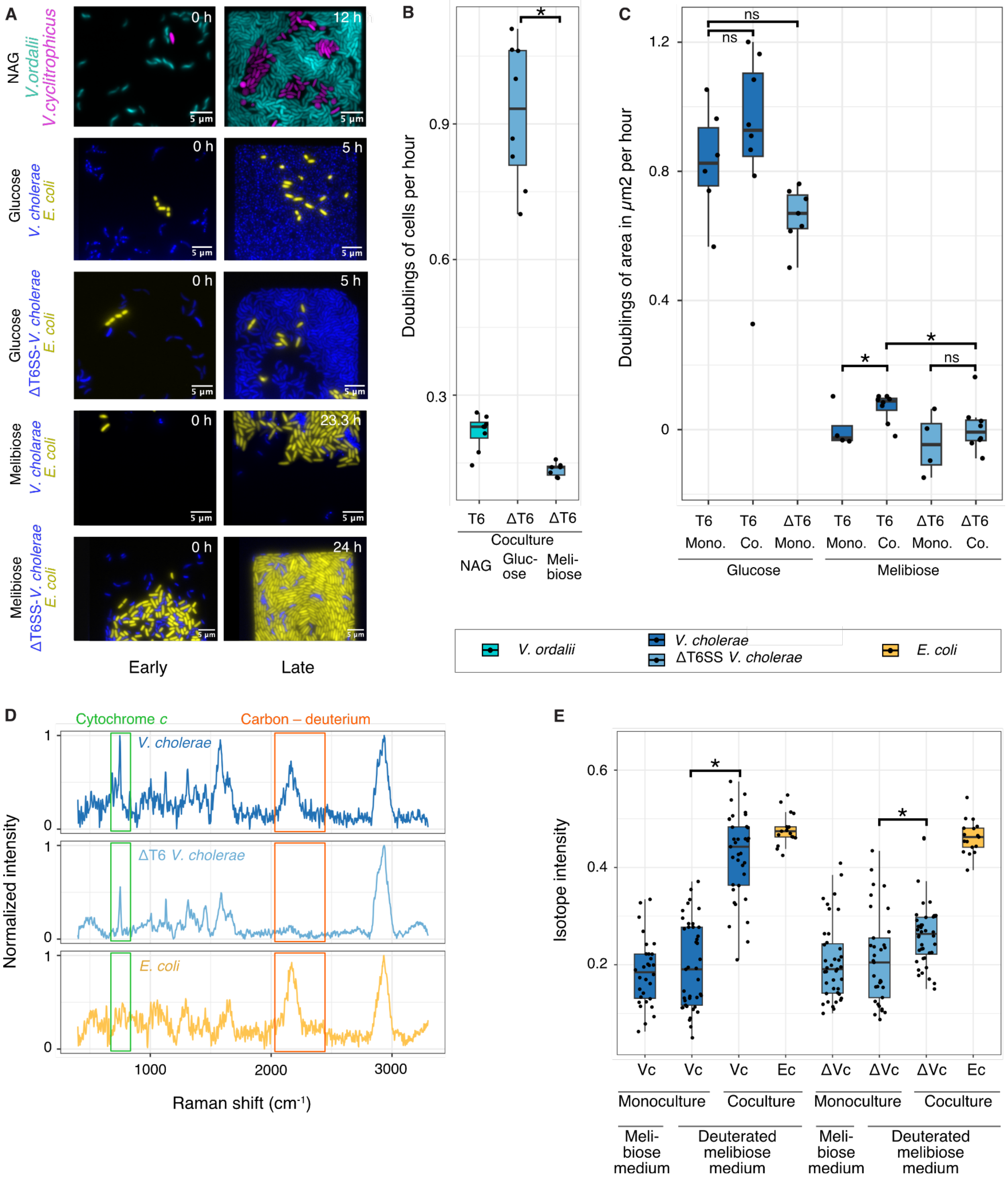


**Fig. S5. Growth benefit and nutrient acquisition through contact-dependent antagonism via the T6SS.**

(A) Representative images of *E. coli* target cells (yellow) and *V. cholerae* cells with and without T6SS (blue) in microfluidic chambers supplied with different carbon sources. For each experimental condition, an early and a late time point is shown. The late time points differ due to differences in growth dynamics of the particular strains on these carbon sources. Notably, in glucose, no lysis of *E. coli* cells was observed, likely due to the fact that glucose protects *E. coli* MG1655 from T6SS-mediated attacks (*39*). (B) Single-cell growth rates of *V. ordalii* and *V. cholerae* without T6SS in microfluidic chambers supplied with different carbon sources. The growth rate was calculated based on cell counts. Wilcoxon rank-sum exact test, p = 0.0003. (C) Area-based growth rates of *V. cholerae* with and without T6SS. Here, growth rates were estimated based on the increase in area occupied by cells because the segmentation of single cells was not possible in all cases (Wilcoxon rank-sum exact test, p = 0.345, 0.051, 0.024, 0.049, and 0.461, left to right). Chamber 4 of *V. cholerae* monoculture on melibiose was excluded, as this chamber exhibited an increased inflow of T6SS cells during imaging, leading to higher area measurements unrelated to cellular growth. (D) A processed Raman spectrum of a single *V. cholerae*, *V. cholerae* T6SS-deletion, and *E. coli* cell after coculturing *V. cholerae* and deuterium-labeled *E. coli* for 9 h. The molecular composition of the cell can be identified from the individual peaks at different wavenumbers (indicated as Raman shift) that correspond to different molecular bonds. *E. coli* cells and *V. cholerae* cells were distinguished by the absence of a cytochrome *c* peak at 750 $cm^{-1}$ in *E. coli* (green box). Only *V. cholerae* with functional T6SS showed incorporation of deuterium, whereas T6SS-deletion mutants did not (orange box). (E) Stable isotope probing (SIP)-Raman microspectroscopy measurement of the carbon-deuterium bond (C-D) relative to the sum of the carbon-hydrogen (C-H) and C-D bonds (n = 30, 41, 35, 17, 39, 33, 40, 17; Wilcoxon rank-sum tests, $p = 9 \times 10^{-12}$ and 0.005, left to right). Vc, ΔVc, and Ec indicate *V. cholerae*, *V. cholerae* T6SS-deletion, and *E. coli*, respectively. Boxplots in (B), (C), and (E) have a central line representing the median, box top and bottom representing the first and third quartiles, respectively, and whiskers extending to the smallest and largest values within 1.5 times the interquartile range from the first and third quartiles.

528

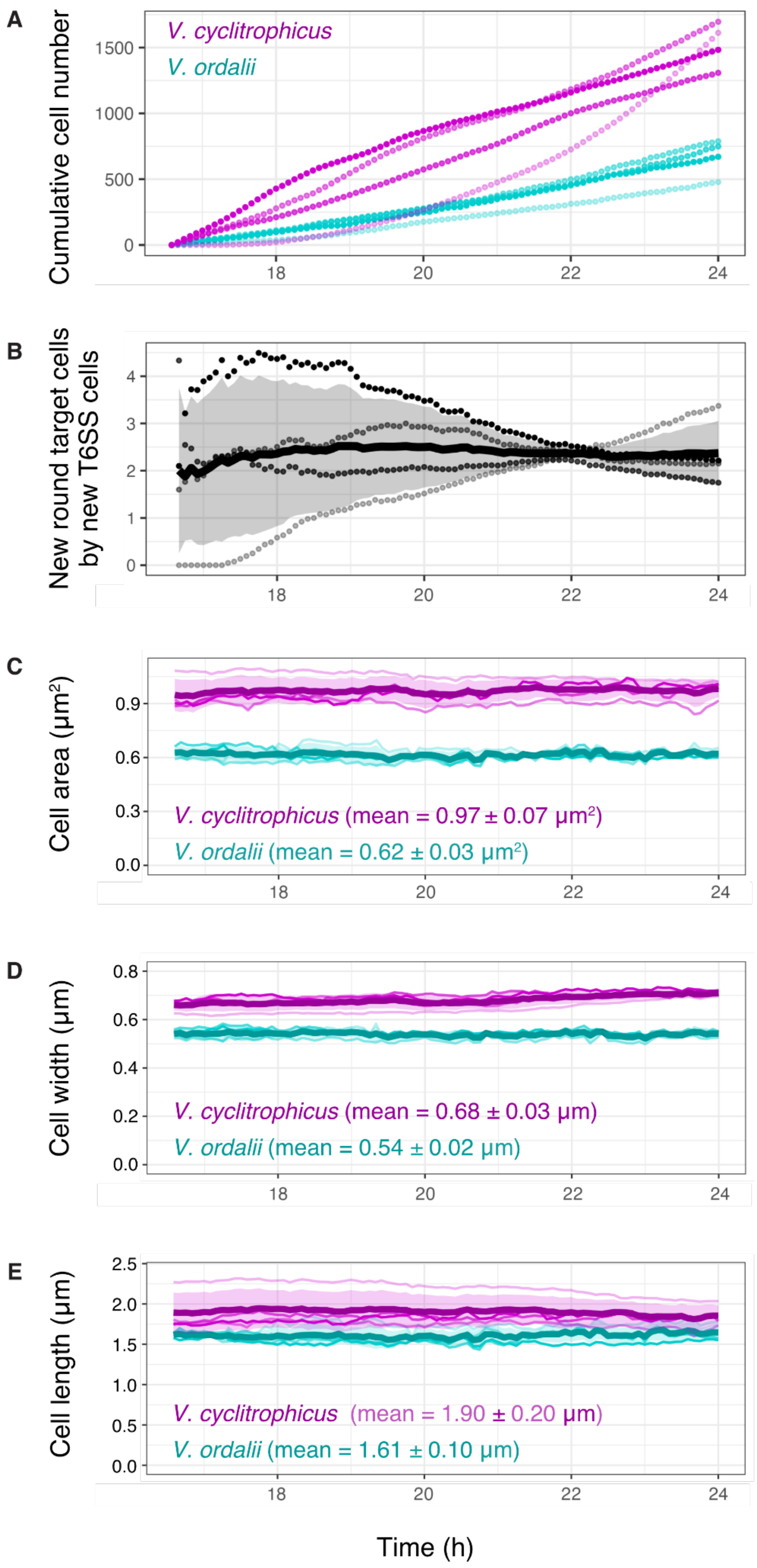

A
Cumulative cell number
V. cyclitrophicus
V. ordalii
B
New round target cells by new T6SS cells
C
Cell area (μm²)
V. cyclitrophicus (mean = 0.97 ± 0.07 μm²)
V. ordalii (mean = 0.62 ± 0.03 μm²)
D
Cell width (μm)
V. cyclitrophicus (mean = 0.68 ± 0.03 μm)
V. ordalii (mean = 0.54 ± 0.02 μm)
E
Cell length (μm)
V. cyclitrophicus (mean = 1.90 ± 0.20 μm)
V. ordalii (mean = 1.61 ± 0.10 μm)
Time (h)

**Fig. S6. Transfer efficiency of biomass from lysing target cells to T6SS cells.** (A) Cumulative number of new round *V. ordalii* target cells (magenta) and new *V. cyclitrophicus* T6SS cells (cyan) in microfluidic chambers starting at 16.75 hours into the experiment. Chambers with few killing events, i.e., a cumulative count of less than 200 round target cells after 24, were excluded from this analysis. (B) Ratio of the cumulative number of new round target cells and new T6SS cells. (C) The average size in terms of area of T6SS cells (cyan) is smaller than that of target cells (magenta). (D) The average cell width of T6SS cells (cyan) is smaller than that of target cells (magenta). It was used to estimate the cell volume of target and T6SS cells. (E) The average cell length of T6SS cells (cyan) is smaller than that of target cells (magenta). It was used to estimate the cell volume of target and T6SS cells. For (A)-(E), different color intensities correspond to different microfluidic chambers, n = 4 chambers, thick line represents the mean of all chambers, shaded background represents the 95% confidence interval.

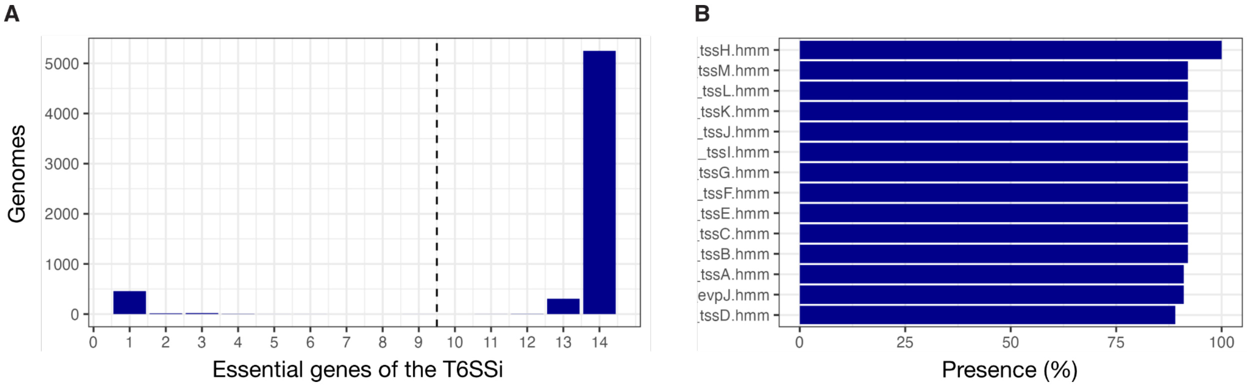


**Fig. S7. Presence of T6SS genes across *Vibrio* genomes.**
(A) Number of essential T6SS subtype i (T6SSi) genes in 6,062 *Vibrio* high-quality genomes from RefSeq. Subtype i is the subtype of the T6SS that is found in proteobacteria such as *Vibrio* (*40*). (B) Presence of each essential gene of the T6SSi in the 6,062 *Vibrio* genomes with any T6SS genes.

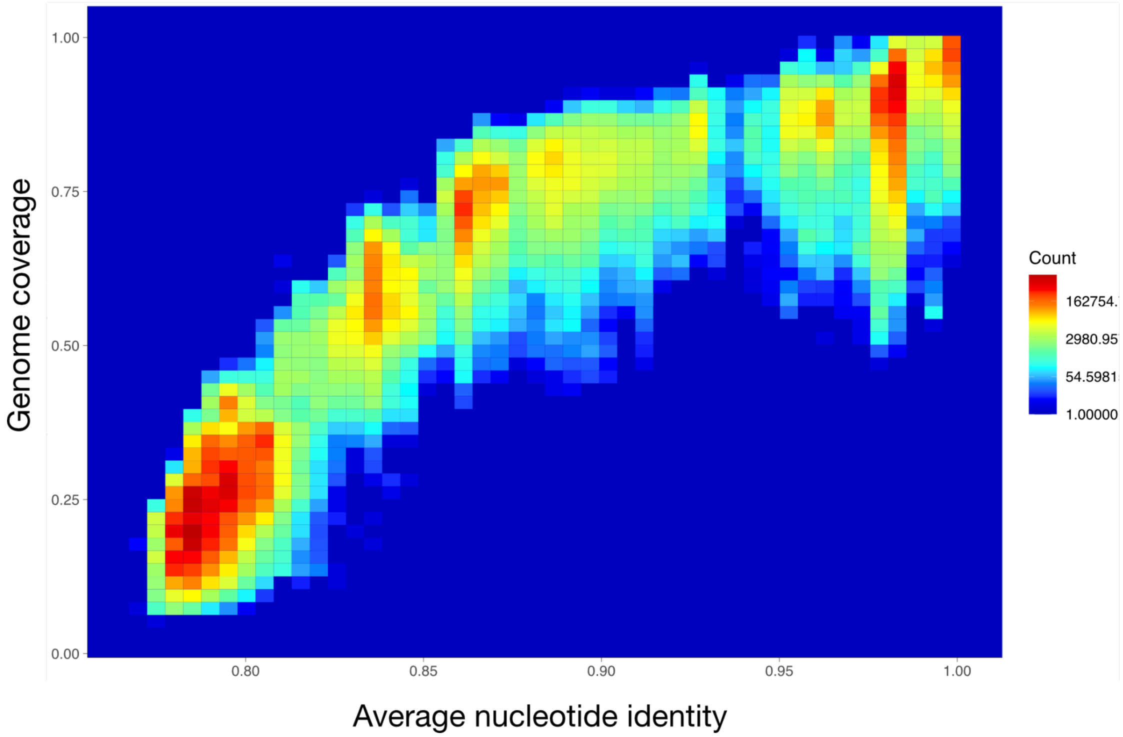


**Fig. S8. Average nucleotide identity among *Vibrio* genomes.**
The average nucleotide identity (ANI) and genome alignment percentage (genome coverage, vertical axis) were computed for pairwise comparisons of 6,062 *Vibrio* genomes and depicted as heatmap. Higher-intensity colors represent a higher density of pairwise comparisons with that particular ANI and genome coverage. Due to the gap in ANI values around 94%, we chose a threshold of 94% ANI to de-replicated the *Vibrio* genomes into OTUs in the following analysis.

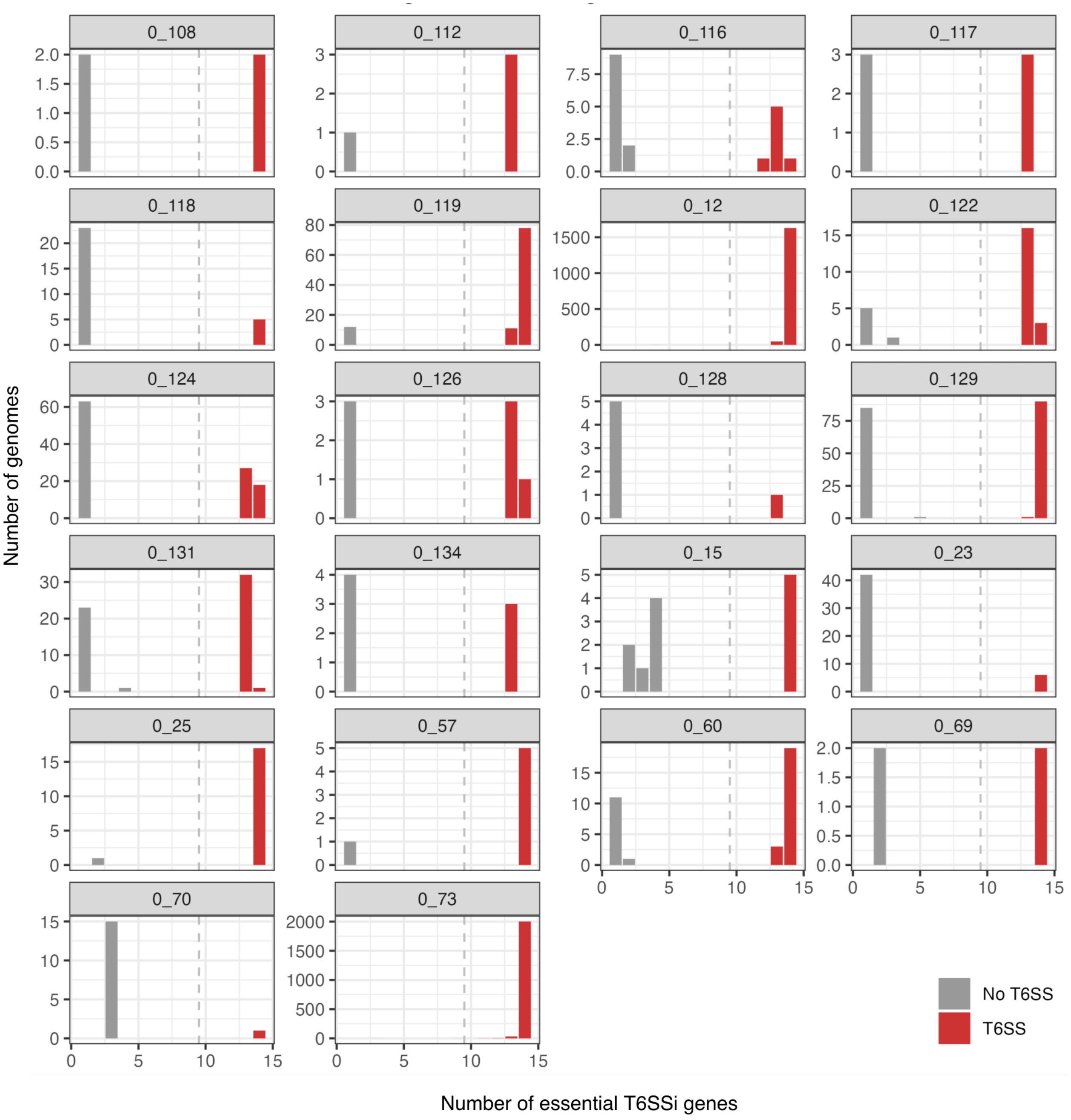


**Fig. S9. Presence of T6SS genes in the genomes of the 94% ANI clusters that contain both *Vibrio* genomes with and without a T6SS.**

The clusters 0_12 and 0_73 were removed from the T6SS-heterogeneous clusters and considered clusters of T6SS-encoding genomes (see Materials and Methods) leaving 20 genome clusters with strain-specific presence of a T6SS.

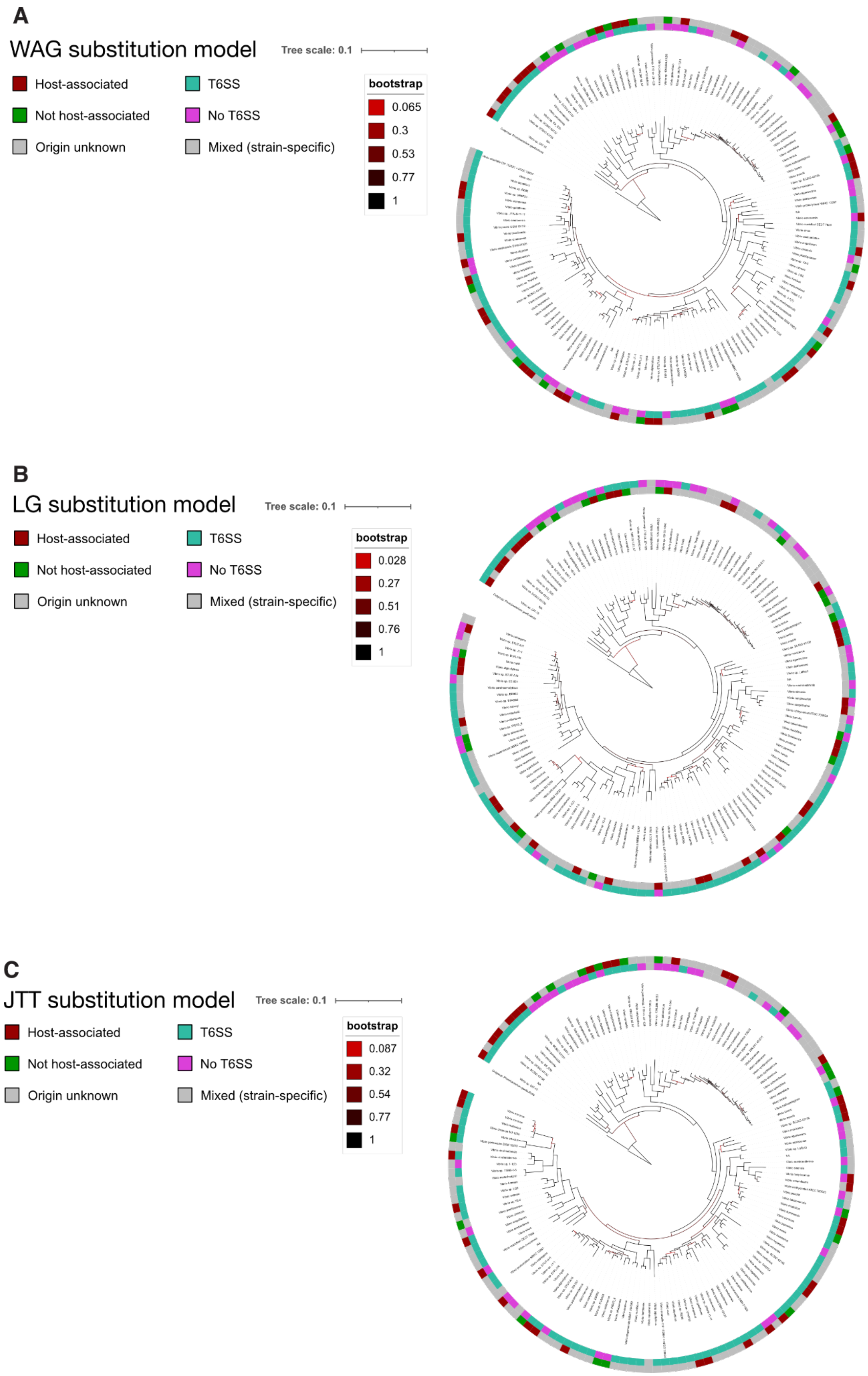
A
WAG substitution model
Tree scale: 0.1
Host-associated
Not host-associated
Origin unknown
T6SS
No T6SS
Mixed (strain-specific)
bootstrap
0.065
0.3
0.53
0.77
1
B
LG substitution model
Tree scale: 0.1
Host-associated
Not host-associated
Origin unknown
T6SS
No T6SS
Mixed (strain-specific)
bootstrap
0.028
0.27
0.51
0.76
1
C
JTT substitution model
Tree scale: 0.1
Host-associated
Not host-associated
Origin unknown
T6SS
No T6SS
Mixed (strain-specific)
bootstrap
0.087
0.32
0.54
0.77
1

**Fig. S10. Phylogenetic tree of 141 representative *Vibrio* genomes.**

The phylogenetic tree was created with GTDBtk, which relies on 120 bacterial single copy marker genes, with *Photobacterium gaetbulicola* as an outgroup. Here, the (A) Whelan and Goldman (WAG), (B) Le and Gascuel (LG), and (C) JTT Jones-Taylor-Thornton (JTT) protein substitution model was used to infer the phylogenetic tree. Bootstrap values displayed as red to black paths. Taxonomic names assigned by NCBI (leaf label). The T6SS label (inner ring) denotes the presence of the T6SS genomes of the OTU (cyan), absence of the T6SS in all genomes of the OTU (magenta), or both presence and absence of the T6SS in genomes of the OTU, indicating strain-specific presence or absence of the T6SS (gray). The origin label (outer ring) states the origin of the isolate as host-associated (red), not host-associated (green), or not recorded on NCBI (gray).

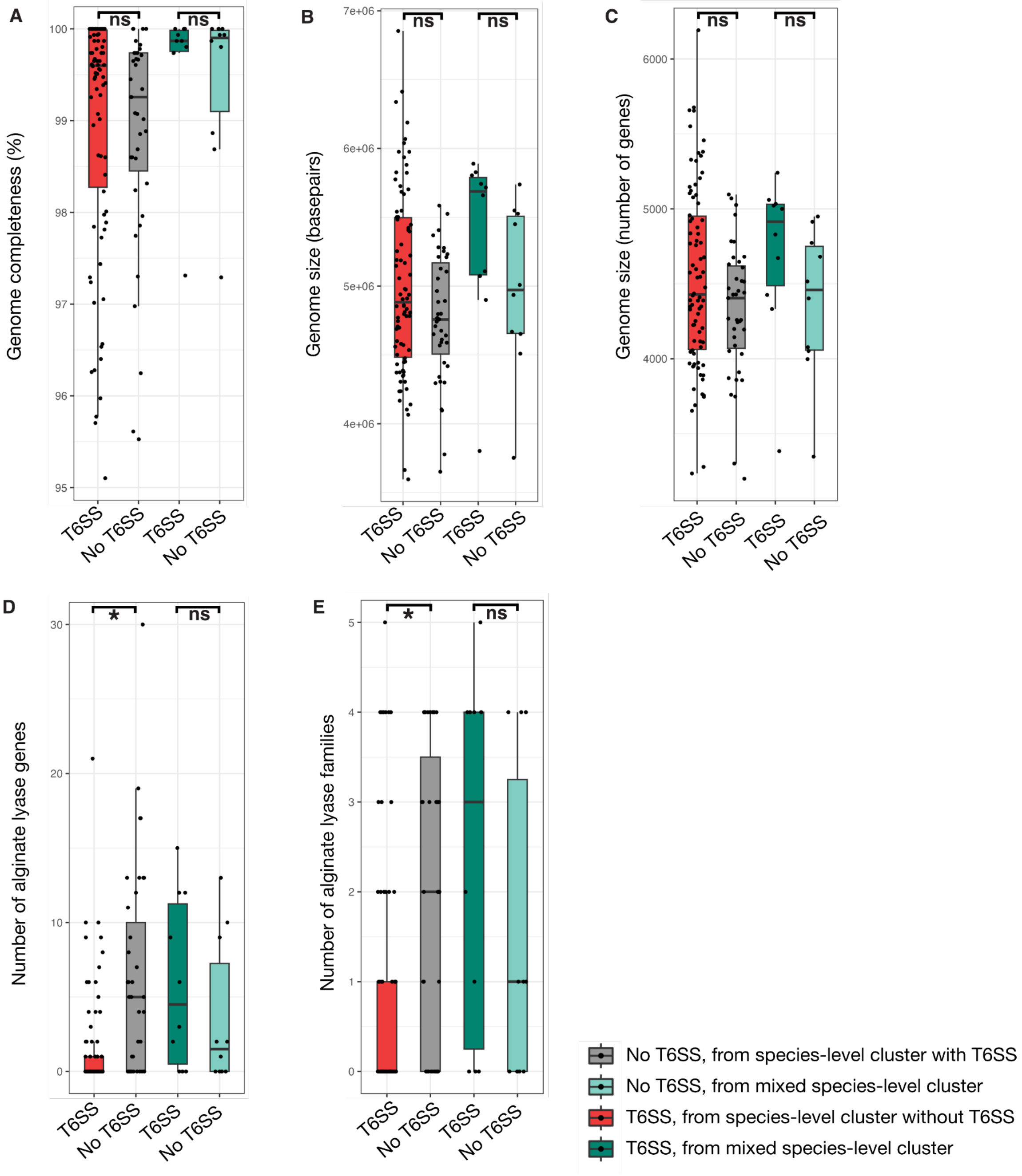


**Fig. S11. Genomic comparison of T6SS-encoding and T6SS-lacking genomes without correction for phylogenetic relationships.**

(A) Genome completeness estimated by checkM for each genome. (B) Genomes size in base pairs of each genome. (C) Genome size in the number of genes of each genome. (D) Number of genes annotated as alginate lyases in each genome. (E) Number of unique alginate lyase families encoded in each genome. Boxplot with central line representing the median, box top and bottom representing the first and third quartiles, respectively, and whiskers extending to the smallest and largest values within 1.5 times the interquartile range from the first and third quartiles.

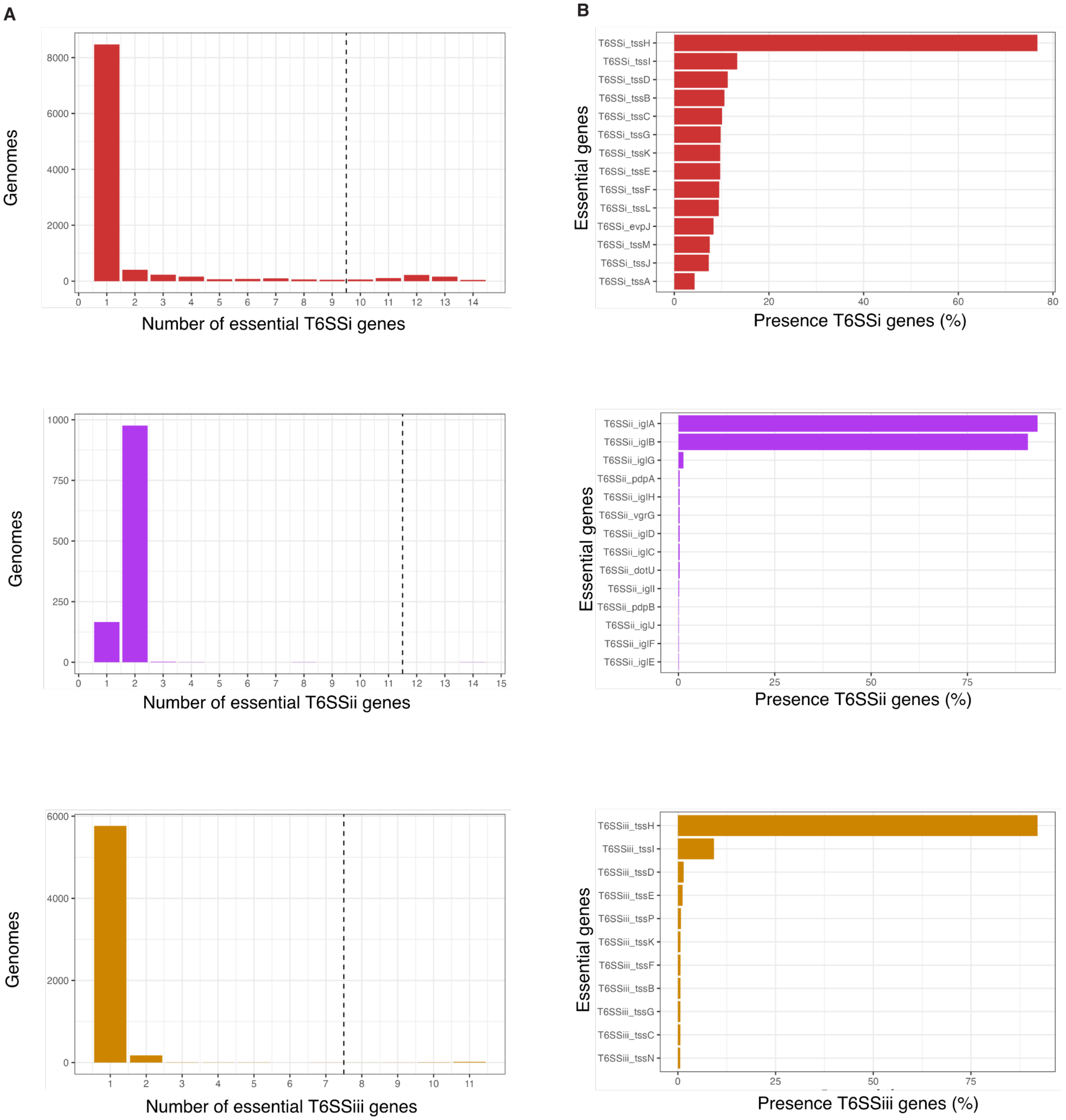


**Fig. S12. Presence of T6SS genes across the ~35,000 genomes of the OMD.**
(A) Number of essential T6SSi genes (top), T6SSii genes (middle), and T6SSiii genes (bottom) in the genomes contained in the OMD. (B) Presence of each essential gene of the T6SSi (top), T6SSiii (middle), and T6SSiii (bottom) in the genomes of the OMD that contained any T6SS genes.

**Table S1. Details and genetic modifications of the studied strains.**

| Organism | Genotype | Plasmid/Features | References |
|---|---|---|---|
| *Vibrio cyclitrophicus* ZF270 | ZF270 wild type | pLL103-GFP, $Chl^R$ *GFP* labeled prey cells | This study |
| *Vibrio ordali* FS144 | FS144 wild type | pVSV208-dsRed, $Chl^R$ *dsRed* labeled killer cells | This study |
| *Vibrio ordali* 12B09-HW44 | 12B09 *Δnrp* | pVSV208-dsRed, $Chl^R$ *dsRed* labeled killer cells | (*3*) This study |
| *Escherichia coli* TB204 | MG1655 *att*P21::*PR-sfGFP* | *sfGFP* labeled prey cells | (*4*) |
| *Escherichia coli* TB205 | MG1655 *att*P21::*PR-mCherry* | *mCherry* labeled prey cells | (*4*) |
| *Vibrio cholerae* 2740-80 B625 | lacZ- , $Str^r$ , *vipA-msfGFP* | C-terminal chromosomal fusion of *msfGFP* to *vipA* | (*41*) |
| *Vibrio cholerae* 2740-80 JS93 | lacZ- , $Str^r$ , *vipA-mCherry2* | C-terminal chromosomal fusion of *mCherry2* to *vipA* | (*41*) |
| *Vibrio cholerae* 2740-80 JS107 | lacZ- , $Str^r$ , *vipA-msfGFP, Δhcp1, Δhcp2* | T6SS sheath deletion in a *vipA-msfGFP* background | (*41*) |
| *Vibrio cholerae* 2740-80 AV009 | lacZ- , $Str^r$ , *vipA-mCherry2, Δhcp1, Δhcp2* | T6SS sheath deletion in a *vipA-mCherry2* background | (*7*) |

**Table S2. Screen for T6SS effector proteins in *V. ordalii* and *V. cholerae.***

A genomic screen with the tool SecRet6 v3 (*42*) revealed that *V. ordalii* FS144 and *V. cholerae* 2740-80 encode multiple T6SS effector proteins, including both slow- and fast-acting toxins based on the classification in Table S3.

| **Toxins detected with blastp-based scanning tool SecReT6** | |
|---|---|
| ***V. ordalii*** | ***V. cholerae*** |
| Tde (hydrolysis nucleic acids) | Tae (degrades peptidoglycan) |
| Tle (degrade membrane phospholipids) | Tde (hydrolysis nucleic acids) |
| Tme (form pores in the inner membrane) | Tle (degrade membrane phospholipids) |
| Tse (unspecified) | Tme (form pores in the inner membrane) |
| | Tse (unspecified) |

**Table S3. Classification of T6SS effector proteins into fast and slow acting toxins.** Classification according to Smith et al. (*43*).

| | **Classification based on Smith *et al.* (2020)** | |
|---|---|---|
| | **Fast lysis** | **Slow lysis** |
| **Toxin target** | Peptidoglycan and phospholipids | DNA, membrane pore-forming, and NAD(P+) |
| **Toxin** | Am-1-3, Mur, Glc, PLA1-2, PLD | Tox43, Tse4, VasX, Tox46 |

**Table S4. Physiological values of the intracellular concentration $C_i$ and half-saturation constant $K_d$.**

| Variable | Measurement | Value | Reference |
|---|---|---|---|
| $K_d$ | Half-saturation constant for amino acid transport systems in *E. coli* at 37 ºC | 1 − 10 μM | (*15*) |
| $C_i$ | Intracellular amino acid levels in *E. coli* in mid-exponential growth phase (OD650 = 0.6) | 1,000 − 60,000 μM | (*14*) |

**Table S5. Distribution of the T6SS in *Vibrio* isolates with host-associated and not host-associated origin.**

The fraction of T6SS-containing genomes was calculated as T6SS-encoding / (T6SS-encoding + T6SS-lacking), excluding the counts of mixed genome clusters that contain both genomes with and without T6SS.

| Environment | T6SS | No T6SS | Mixed | SUM | Fraction of T6SS (%) |
|---|---|---|---|---|---|
| **NA** | 49 | 21 | 15 | 85 | 70.0 |
| **Host-associated** | 28 | 7 | 3 | 38 | 80.0 |
| **Not host-associated** | 5 | 11 | 2 | 18 | 31.3 |
| **SUM** | 82 | 39 | 20 | 141 | 67.8 |

**Table S6. Distribution of the T6SS in Vibrio isolates with host-associated and different environmental origins.**
The fraction of T6SS was calculated as T6SS-encoding / (T6SS-encoding + T6SS-lacking), excluding the counts of mixed genome clusters that contain both genomes with and without T6SS.

| Environment | T6SS | No T6SS | Mixed | SUM | Fraction of T6SS (%) |
|---|---|---|---|---|---|
| **NA** | 49 | 21 | 15 | 85 | 70.0 |
| **Host-associated** | 28 | 7 | 3 | 38 | 80.0 |
| **Marine** | 2 | 7 | 1 | 10 | 22.2 |
| **Marine water** | 2 | 2 | 0 | 4 | 50.0 |
| **Rice root** | 1 | 0 | 1 | 2 | 100.0 |
| **Coral reef** | 0 | 1 | 0 | 1 | 0.0 |
| **Marine sediment** | 0 | 1 | 0 | 1 | 0.0 |
| **SUM** | 82 | 39 | 20 | 141 | 67.8 |

**Movies**

**Movie 1. Time-lapse video of a *Vibrio ordalii* monoculture (T6SS cells) within a representative microfluidics chamber fed with 0.1% *N*-acetylglucosamine (GlcNAc) as the sole carbon source.** Images were captured every 10 min. Cells are false-colored in cyan based on the fluorescence signal of the dsRed plasmid-based marker. The scale bar corresponds to 5 µm. Image corresponds to pos6 from the raw dataset.

**Movie 2. Time-lapse video of a *Vibrio ordalii* monoculture (T6SS cells) within a representative microfluidics chamber fed with 0.1% alginate as the sole carbon source.** Images were captured every 10 min. Cells are false-colored in cyan based on the fluorescence signal of the dsRed plasmid-based marker. The scale bar corresponds to 5 µm. Image corresponds to pos8 from the raw dataset.

**Movie 3. Time-lapse video of Vibrio *cyclitrophicus* (target cells) and Vibrio *ordalii* (T6SS cells) within a representative microfluidics chamber fed with 0.1% alginate as the sole carbon source.** Images were captured every 5 min. Cells are false-colored in magenta (target cells) or cyan (T6SS cells) based on fluorescence signals of the GFP and dsRed plasmid-based markers, respectively. The scale bar corresponds to 5 µm. Image corresponds to pos12 from the raw dataset.

**Movie 4. Time-lapse video of a *Vibrio cholerae* monoculture (T6SS cells) within a representative microfluidics chamber fed with 0.1% melibiose as the sole carbon source.** Images were captured every 5 min. Cells are false-colored in blue based on the fluorescence signal of the msfGFP chromosomal marker. The scale bar corresponds to 10 µm. Image corresponds to pos16 from the raw dataset.

**Movie 5. Time-lapse video of Escherichia *coli* (target cells) and *Vibrio cholerae* (T6SS cells) within a representative microfluidics chamber fed with 0.1% melibiose as the sole carbon source.** Images were captured every 5 min. Cells are false-colored in yellow (target cells) or blue (T6SS cells) based on fluorescence signals of the mCherry2 and sfGFP chromosomal markers, respectively. The scale bar corresponds to 10 µm. Image corresponds to pos5 from the raw dataset.

**Movie 6. Time-lapse video of *Escherichia coli* (target cells) and T6SS-deletion mutant *Vibrio cholerae* cells within a representative microfluidics chamber fed with 0.1% melibiose as the sole carbon source.** Images were captured every 5 min. Cells are false-colored in yellow (target cells) or blue (T6SS-deletion mutant cells) based on fluorescence signals of the sfGFP and mCherry2 chromosomal markers, respectively. The scale bar corresponds to 5 µm. Image corresponds to pos15 from the raw dataset.

M. DeAngelis, V. Denef, S. E. Denman, A. Desta, H. Dionisi, J. Dodsworth, N. Dombrowski, T. Donohue, M. Dopson, T. Driscoll, P. Dunfield, C. L. Dupont, K. A. Dynarski, V. Edgcomb, E. A. Edwards, M. S. Elshahed, I. Figueroa, B. Flood, N. Fortney, C. S. Fortunato, C. Francis, C. M. M. Gachon, S. L. Garcia, M. C. Gazitua, T. Gentry, L. Gerwick, J. Gharechahi, P. Girguis, J. Gladden, M. Gradoville, S. E. Grasby, K. Gravuer, C. L. Grettenberger, R. J. Gruninger, J. Guo, M. Y. Habteselassie, S. J. Hallam, R. Hatzenpichler, B. Hausmann, T. C. Hazen, B. Hedlund, C. Henny, L. Herfort, M. Hernandez, O. S. Hershey, M. Hess, E. B. Hollister, L. A. Hug, D. Hunt, J. Jansson, J. Jarett, V. V. Kadnikov, C. Kelly, R. Kelly, W. Kelly, C. A. Kerfeld, J. Kimbrel, J. L. Klassen, K. T. Konstantinidis, L. L. Lee, W.-J. Li, A. J. Loder, A. Loy, M. Lozada, B. MacGregor, C. Magnabosco, A. Maria Da Silva, R. M. McKay, K. McMahon, C. S. McSweeney, M. Medina, L. Meredith, J. Mizzi, T. Mock, L. Momper, M. A. Moran, C. Morgan-Lang, D. Moser, G. Muyzer, D. Myrold, M. Nash, C. L. Nesbø, A. P. Neumann, R. B. Neumann, D. Noguera, T. Northen, J. Norton, B. Nowinski, K. Nüsslein, M. A. O’Malley, R. S. Oliveira, V. Maia De Oliveira, T. Onstott, J. Osvatic, Y. Ouyang, M. Pachiadaki, J. Parnell, L. P. Partida-Martinez, K. G. Peay, D. Pelletier, X. Peng, M. Pester, J. Pett-Ridge, S. Peura, P. Pjevac, A. M. Plominsky, A. Poehlein, P. B. Pope, N. Ravin, M. C. Redmond, R. Reiss, V. Rich, C. Rinke, J. L. M. Rodrigues, W. Rodriguez-Reillo, K. Rossmassler, J. Sackett, G. H. Salekdeh, S. Saleska, M. Scarborough, D. Schachtman, C. W. Schadt, M. Schrenk, A. Sczyrba, A. Sengupta, J. C. Setubal, A. Shade, C. Sharp, D. H. Sherman, O. V. Shubenkova, I. N. Sierra-Garcia, R. Simister, H. Simon, S. Sjöling, J. Slonczewski, R. S. Correa De Souza, J. R. Spear, J. C. Stegen, R. Stepanauskas, F. Stewart, G. Suen, M. Sullivan, D. Sumner, B. K. Swan, W. Swingley, J. Tarn, G. T. Taylor, H. Teeling, M. Tekere, A. Teske, T. Thomas, C. Thrash, J. Tiedje, C. S. Ting, B. Tully, G. Tyson, O. Ulloa, D. L. Valentine, M. W. Van Goethem, J. VanderGheynst, T. J. Verbeke, J. Vollmers, A. Vuillemin, N. B. Waldo, D. A. Walsh, B. C. Weimer, T. Whitman, P. Van Der Wielen, M. Wilkins, T. J. Williams, B. Woodcroft, J. Woolet, K. Wrighton, J. Ye, E. B. Young, N. H. Youssef, F. B. Yu, T. I. Zemskaya, R. Ziels, T. Woyke, N. J. Mouncey, N. N. Ivanova, N. C. Kyrpides, E. A. Eloe-Fadrosh, A genomic catalog of Earth’s microbiomes. *Nat Biotechnol* **39**, 499–509 (2021).

38. A. K. Stubbusch, J. M. Keegstra, J. Schwartzman, S. Pontrelli, E. E. Clerc, R. Stocker, C. Magnabosco, O. T. Schubert, M. Ackermann, G. G. D’Souza, Polysaccharide breakdown products drive degradation-dispersal cycles of foraging bacteria through changes in metabolism and motility. *eLife* **13** (2024).
39. C. V. Crisan, H. L. Nichols, S. Wiesenfeld, G. Steinbach, P. J. Yunker, B. K. Hammer, Glucose confers protection to Escherichia coli against contact killing by Vibrio cholerae. *Scientific Reports* **11**, 1–11 (2021).
40. R. Gallegos-Monterrosa, S. J. Coulthurst, The ecological impact of a bacterial weapon: microbial interactions and the Type VI secretion system. *FEMS Microbiology Reviews* **45**, fuab033 (2021).
41. A. Vettiger, M. Basler, Type VI Secretion System Substrates Are Transferred and Reused among Sister Cells. *Cell* **167**, 99-110.e12 (2016).
42. J. Zhang, J. Guan, M. Wang, G. Li, M. Djordjevic, C. Tai, H. Wang, Z. Deng, Z. Chen, H.-Y. Ou, SecReT6 update: a comprehensive resource of bacterial Type VI Secretion Systems. *Sci. China Life Sci.* **66**, 626–634 (2023).
43. W. P. J. Smith, A. Vettiger, J. Winter, T. Ryser, L. E. Comstock, M. Basler, K. R. Foster,

The evolution of the type VI secretion system as a disintegration weapon. *PLoS Biology* **18**, 1–26 (2020).
